# Accurate model and ensemble refinement using cryo-electron microscopy maps and Bayesian inference

**DOI:** 10.1101/2023.10.18.562710

**Authors:** S. E. Hoff, F. E. Thomasen, K. Lindorff-Larsen, M. Bonomi

## Abstract

Converting cryo-electron microscopy (cryo-EM) data into high-quality structural models is a challenging problem of outstanding importance. Current refinement methods often generate unbalanced models in which physico-chemical quality is sacrificed for excellent fit to the data. Furthermore, these techniques struggle to represent the conformational heterogeneity averaged out in low-resolution regions of density maps. Here we introduce EMMIVox, a Bayesian inference approach to determine single-structure models as well as structural ensembles from cryo-EM maps. EMMIVox automatically balances experimental information with accurate physico-chemical models of the system and the surrounding environment, including waters, lipids, and ions. Explicit treatment of data correlation and noise as well as inference of accurate B-factors enable determination of structural models and ensembles with both excellent fit to the data and high stereochemical quality, thus outperforming state-of-the-art refinement techniques. EMMIVox represents a flexible approach to determine high-quality structural models that will contribute to advancing our understanding of the molecular mechanisms underlying biological functions.

## Introduction

Cryo-electron microscopy (cryo-EM) has become a powerful tool to determine the structures of complex biological systems. Recent advances in physical instrumentation and image processing algorithms have pushed the boundaries of what can be achieved with this technique in terms of resolution and coverage across a wide spectrum of system size and complexity^1^. While most single-particle cryo-EM density maps still have a resolution between 3 Å and 4 Å (Fig. S1A), the number of systems determined at atomic resolution is steadily increasing, with the current record set by Apoferritin, resolved in 2020 at ∼1.2 Å^2, 3^. At the same time, tremendous progress in cryo-electron tomography (cryo-ET) is enabling the determination of the structure of complex biological systems *in situ* at sub-nanometer resolution^4^ (Fig. S1B). It is also becoming increasingly clear that single-structure models are not always a faithful representation of the three-dimensional (3D) cryo-EM density maps^5^. Small-scale continuous dynamics of functionally important flexible regions within biomolecules are typically averaged out during the reconstruction of 3D maps resulting in fuzzy density regions with reduced resolution. To accurately interpret these regions, we should therefore move away from single-structure models towards ensembles of conformations^6, 7^. In all these situations, it is of paramount importance to convert cryo-EM data into high-quality structural models, for example for *in silico* structure-based drug design^8^ or training deep learning approaches on accurate structural data^9^.

Over the years, a variety of different metrics have been proposed to evaluate the quality of structural models obtained from cryo-EM maps^10-13^. Generally speaking, these metrics fall into two categories: how well a model (or set of models) explains the observed map or directly the single-particle images (*fit to the data*) and how good the model is in terms of basic stereochemical parameters, such as length of chemical bonds, backbone and sidechain dihedral distributions, and clashes between close atoms. To evaluate the fit to the data, the most common approach is to compare the experimental map with the map predicted from a model, for example by calculating the cross correlation between the two maps over the entire 3D space or in the proximity of the structural model. In low-resolution areas, the density can be predicted either as an average over multiple conformations or from a single-structure model by introducing temperature factors (*B-factors*), which implicitly account for intrinsic dynamics and other reasons behind the observed fuzzy density, such as errors in the image alignment or local damage caused by the electron beam. The main challenge in determining a high-quality structural model is to obtain a good balance between the fit to the observed density map and the overall stereochemical quality of the model.

Several modelling approaches have been developed to refine structural models into cryo-EM maps^14^. These techniques rely on various approaches including homology modelling^15^, rigid-body fitting^16-19^, flexible refinement^20-28^, integrative modelling^29, 30^, and machine learning to automate model fitting to 3D density maps^31^. Most of these methods do not optimize B-factors, and therefore, while they are extremely useful to create structural models that occupy the space defined by the cryo-EM density, a quantitative evaluation of the model fit to the experimental map with the metrics commonly used for PDB validation is challenging. On the other hand, a handful of methods have been proposed to model structural ensembles from either 2D single-particle images^32-34^ or 3D maps^22, 35, 36^. As of today, one of the most popular refinement software commonly used prior to depositing models in the PDB database is PHENIX^23^, which enables real-space refinement, optimization of B-factors, and modelling residues in alternative conformations. The approach implemented in PHENIX relies on an empirical scoring function to maximize the correlation between the cryo-EM map predicted from a model and the experimental map while trying to preserve stereochemical properties. This approach typically leads to an excellent fit to the data (Fig. S2AB) but often at the expense of physico-chemical properties, especially in terms of clashes between atoms (Fig. S2CD).

Here we present EMMIVox, a computational approach to determine single-structure models as well as conformational ensembles using cryo-EM maps. EMMIVox is based on a Bayesian inference framework^37^ to balance automatically the experimental information with state-of-the-art physico-chemical models of the system and the surrounding environment. Explicit treatment of data correlation and uncertainty as well as accurate inference of B-factors contribute to the determination of structural models with excellent fit to the data without sacrificing the stereochemical quality of the models. We benchmarked our approach on nine complex biological systems and demonstrated that EMMIVox models outperformed those obtained with state-of-the-art refinement techniques and deposited in the PDB database in terms of several quality metrics. We also illustrate how EMMIVox can be used in combination with medium-low resolution cryo-EM maps to refine coarse-grained models of large protein complexes and to determine conformational ensembles describing the structural heterogeneity hidden in low-resolution areas of atomistic cryo-EM maps. EMMIVox is implemented in the open-source, freely available PLUMED library^38, 39^ (www.plumed.org) and aims at setting a new standard for single-structure and ensemble refinement by optimally integrating cryo-EM maps with accurate atomistic and coarse-grained physico-chemical models of the system as well as other experimental data, when available.

## Results

This section is organized as follows. We first provide a general overview of the EMMIVox approach and illustrate its accuracy in refining both atomistic and coarse-grained single-structure models using cryo-EM maps at various resolutions. We then focus on ensemble refinement and present the results of our benchmark as well as one case study: the GPT type 1a tau filament from progressive supranuclear palsy neurodegenerative disease^40^.

### Overview of the EMMIVox approach

EMMIVox is based on a Bayesian inference framework to generate a *hybrid energy function* that combines the molecular mechanics force fields used in classical Molecular Dynamics (MD) simulations with spatial restraints to enforce the agreement of a structural model with the density observed in the voxels of a cryo-EM map. To balance automatically stereochemical quality of the models and fit to the data, EMMIVox: *i)* pre-filters the voxels of a cryo-EM map to reduce the correlation between experimental data points and therefore help avoid data overweighting; *ii)* uses prior models of uncertainty in the experimental data obtained from independent 3D reconstructions (*half maps*) while allowing for the presence of random and systematic errors in the map; and *iii)* builds spatial restraints weighted by the estimated accuracy in each voxel. Furthermore, EMMIVox exploits modern atomistic and coarse-grained force fields to accurately describe the physico-chemical properties of a biological system as well as its environment, including water molecules, ions, lipids, and small molecules. A Monte Carlo optimization of residue-level B-factors coupled with structural refinement guided by the EMMIVox hybrid energy function enables the determination of single-structure models that fit the observed cryo-EM map while preserving a high stereochemical quality. Instead of implicitly modelling local dynamics with B-factors, EMMIVox can be combined with metainference^41^ to obtain structural ensembles that explicitly represent the continuous dynamics of highly flexible regions of the system. A detailed description of EMMIVox is provided in Materials and Methods.

### Accurate atomistic single-structure refinement

We first benchmarked the accuracy of EMMIVox in refining single-structure models using as test systems the GPT type 1a tau filament (1.90 Å, PDB 7p6a)^40^, the ChRmine channelrhodopsin (2.02 Å, PDB 7w9w)^42^, the Anaplastic lymphoma kinase extracellular domain fragment in complex with an activating ligand (2.27 Å, PDB 7n00)^43^, a dimeric unphosporylated Pediculus humanus protein kinase (2.35 Å, PDB 7t4n)^44^, the human CDK-activating kinase bound to the inhibitor ICEC0942 (2.50 Å, PDB 7b5o)^45^, the major facilitator superfamily domain containing 2A in complex with LPC-18:3 (3.03 Å, PDB 7mjs)^46^, the Escherichia coli PBP1b (3.28 Å, PDB 7lq6)^47^, the mammalian peptide transporter PepT2 (3.50 Å, PDB 7nqk)^48^, and the SPP1 bacteriophage tail tube (4.00 Å, PDB 6yeg)^49^. These examples include soluble and membrane proteins, protein complexes, as well as systems containing ordered waters, small molecules, and lipids.

The quality of the EMMIVox models was evaluated and compared to the deposited PDBs using four key metrics. The physico-chemical quality was quantified by the clashscore^50^ and MolProbity score^50^, while the fit to the data using the correlation coefficient between predicted and observed maps in the proximity of the model (*CC*_*mask*_)^51^ and the EMRinger score^52^. EMMIVox models generally showed improved quality in all four metrics (Fig. 1). The improvements are particularly pronounced for clashscore that indicates the absence of serious clashes between atoms in the EMMIVox models, and the MolProbity score, suggesting a superior overall stereochemical quality. At the same time, EMMIVox models often improved the *CC*_*mask*_ indicating a better fit to the experimental cryo-EM map, and the EMRinger scores, particularly in maps with resolution better than 4 Å. Overall, the four metrics indicate that EMMIVox generates single-structure models that improve upon current models deposited in the PDB.

**Fig. 1.**
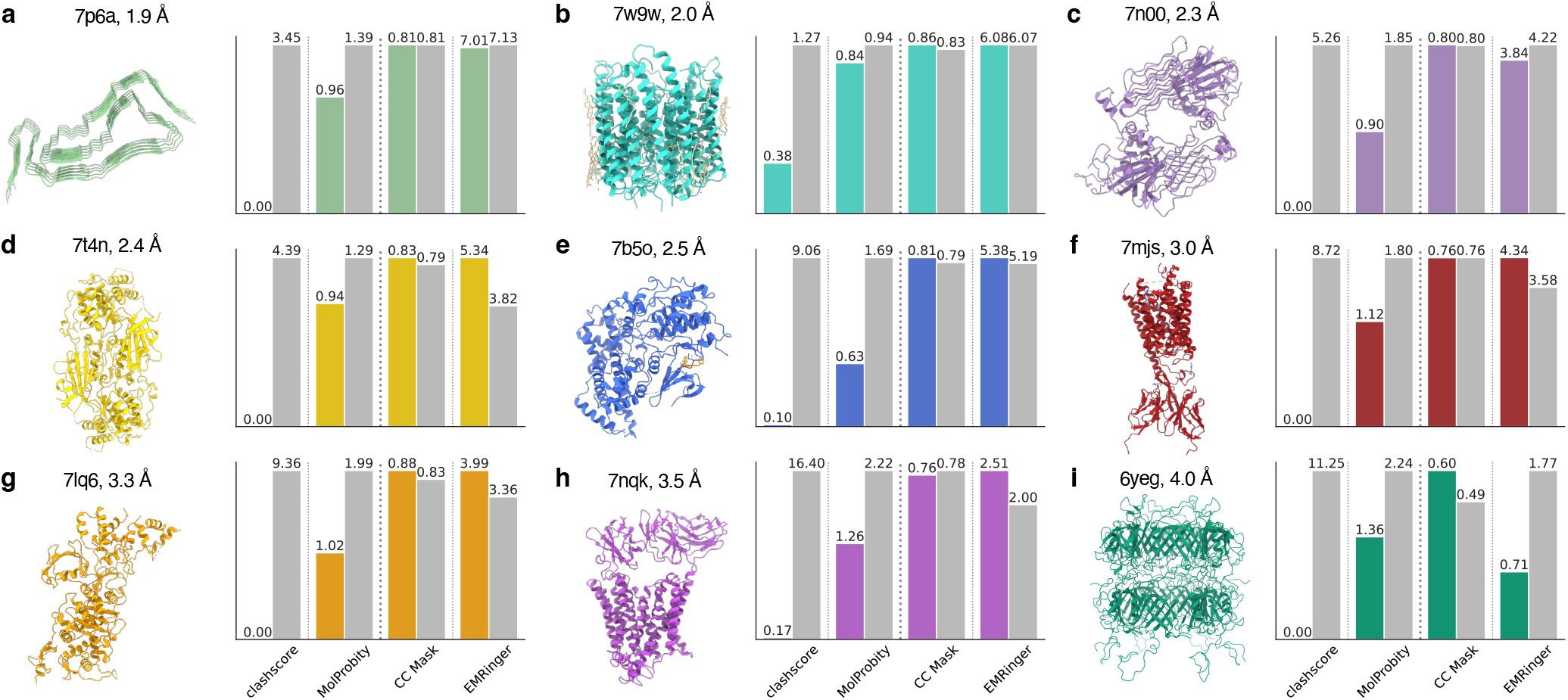
EMMIVox single-structure refinement benchmark. **a-i)** Assessment of the quality of the EMMIVox single-structure model (colored bars) and deposited PDB (grey bars) for each of the nine systems in our benchmark set. Models are evaluated based on their stereochemical quality and fit with the experimental cryo-EM map. The stereochemical quality is measured by two metrics: *i)* clashscore, which corresponds to the number of serious clashes per 1000 atoms, and *ii)* MolProbity score, which is a global measure of quality that combines clashscore, percentage of Ramachandran dihedrals in non-favored regions and percentage of bad sidechain rotamers. For both metrics, lower values correspond to higher quality models. The model fit to the data is measure by *i) CC*_*mask*_, which is the cross-correlation between experimental map and map predicted from the model in the proximity of the structural model, and *ii)* EMRinger, which measures the precise fitting of an atomic model into the map. For both metrics, higher values correspond to models that better fit the data.

In the following sections, we will focus on the different EMMIVox components that contribute to the observed high quality of the models. Three key aspects affect the balance between stereochemical quality and fit to the data: *i)* the number of experimental data points (voxels) that are fit using spatial restraints; *ii)* the strength of these spatial restraints; *iii)* the accuracy of the cryo-EM map predictor from a model.

#### Removal of correlated data leads to balanced model refinement

Neighboring voxels of a cryo-EM map contain correlated information, with the strength of correlation depending on the voxel size and the map resolution. Ignoring such correlation leads to overcounting the number of (independent) data points available and ultimately biasing the refinement towards (over)fitting the data at the expense of the stereochemical quality of the models. To address this point, we developed a pre-filtering procedure to subsample the set of cryo-EM voxels and reduce data correlation (Materials and Methods). We first tested our procedure using over 2400 cryo-EM density maps deposited in the EMDB with resolution ranging from 1.78 Å to 4 Å. Upon removal of correlated data, the median number of voxels per model atom decreased as the resolution of the cryo-EM map worsened and became independent of the voxel size (Fig. S3). This indicates, as expected, that the amount of spatial information provided by high-resolution maps is greater than in medium-low resolution maps.

We then benchmarked the quality of the EMMIVox single-structure refinement as a function of the amount of data removed. Removing a large portion of correlated data led to better stereochemical models compared to utilizing all available voxels (Fig. S4 and S5) at the expense of the quality of the fit to the entire cryo-EM map (Fig. S6 and S7). As fewer points were removed, the balance shifted towards the fit to the data. We therefore identified a correlation coefficient equal to 0.8 as the optimal threshold for data removal in EMMIVox refinement to guarantee a good balance between stereochemical quality of the model and fit to the data. In our benchmark set, this threshold corresponded on average to removing 45% (± 30%) of the voxels in the proximity of the deposited PDB, resulting in 13 ± 5 voxels per atom.

#### Bayesian noise models reduce data overfitting

In EMMIVox, the strength of the spatial restraints used to enforce the model agreement with the cryo-EM map reflects the estimated accuracy (*noise level*) of the density in each voxel. Our Bayesian inference framework enables us to estimate the noise level on the fly based on the consistency between cryo-EM data, physico-chemical prior and potentially additional experimental data (Materials and Methods), and ultimately to downweigh voxels that are considered outlier data points during refinement. To guide noise inference, we developed priors based on the density variability in each voxel calculated from two independent 3D reconstructions, or *half maps*. In regions of the map where large variations are observed, our approach helps to avoid overfitting the data.

To exemplify this point, we examined the relation between per-residue fit to the map (local *CC*_*mask*_) and median noise level of the voxels around a given residue. This analysis illustrates that in the EMMIVox models residues in low-noise regions sometimes fit the map even better than in the deposited PDB, while in regions with high noise level the local fit to the map is often much worse (Fig. 2A and S8). In these regions, during refinement EMMIVox correctly reduced the weight of the experimental data in favor of the molecular mechanics forcefield, ultimately increasing the overall stereochemical quality of the model. To determine which specific physico-chemical properties of the system were better represented in our models, we computed the total number of atom clashes, hydrogen bonds, and salt bridges in the EMMIVox models and deposited PDBs across our entire benchmark (Fig. 2B). EMMIVox dramatically reduced the number of serious clashes observed in the deposited PDBs and optimized the positions of backbone and sidechain atoms so as to better describe hydrogen bond and salt bridge geometries (Fig. 2C). The improvement in these three physico-chemical fingerprints is more accentuated for residues in regions of the cryo-EM map with high noise, supporting the ability of our noise models to shift, when needed, the balance of the refinement towards the accurate molecular mechanics forcefield used by EMMIVox.

**Fig. 2.**
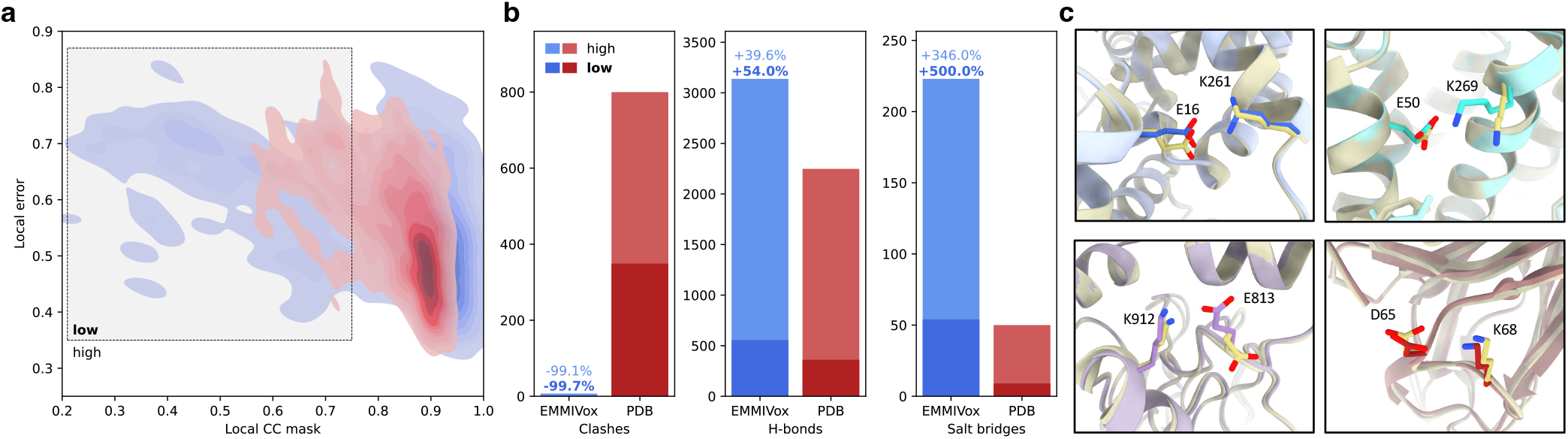
EMMIVox noise models reduce data overfitting and improve stereochemical quality. **a)** Relation between individual residue fit to the experimental cryo-EM map (local *CC*_*mask*_) and local error in the map in the proximity of each residue for the single-structure EMMIVox model (blue) and the deposited PDB (red) in the case of ChRmine channelrhodopsin (2 Å, PDB 7w9w). The local error is calculated from the difference between the two cryo-EM half maps as the median value across all the voxels associated to a residue by Voronoi tessellation. The same analysis for all the other systems of our benchmark set is reported in Fig. S8. **b)** Total number of clashes, hydrogen bonds, and salt bridges in the EMMIVox single-structure models (blue) and deposited PDBs (red) across all the nine systems of our benchmark set. Dark and light bars indicate clashes, hydrogen bonds, and salt bridges that involve residues of the EMMIVox model in low and high *CC*_*mask*_ regions, respectively. Relative variations of these three physico-chemical fingerprints between PDB and EMMIVox models are indicated on top of the blue bars, separately for low and high *CC*_*mask*_ regions. **c)** Examples of salt bridges formed upon EMMIVox refinement of the deposited PDBs (grey).

#### Accurate cryo-EM map prediction with Bayesian inference of B-factors

During refinement and validation, a predictor of cryo-EM density from a 3D model (*forward model*) is required to measure the agreement of a model with the experimental map. Per-residue B-factors need to be determined to smoothen the prediction obtained with classical forward models derived from Gaussian fits of electron scattering factors and thus to explain fuzzy densities with an individual conformation. To reduce data overfitting, PHENIX adds restraints to avoid B-factors of residues close in space to deviate too much from each other. In presence of sharp transitions between ordered and disordered regions, these additional restraints lead to under (over) estimating the B-factors corresponding to flexible (rigid) residues, effectively compressing the space that B-factors can sample. Inspired by PHENIX, we developed a Bayesian inference approach to determine B-factors that allows these restraints to be violated, especially in regions corresponding to order-disorder transitions (Materials and Methods; Fig. S9). The B-factors inferred by EMMIVox contribute to a more accurate density prediction across the whole system and ultimately to increase the model fit to the data.

### Coarse-grained single-structure refinement

For prospective use in single-structure refinement using medium-low resolution cryo-EM and cryo-ET data, we developed a forward model to predict density maps from coarse-grained Martini 3 models, in which each amino acid is represented by a few beads^53^ (Materials and Methods). To evaluate the accuracy of our coarse-grained forward model, we calculated density maps from the Martini representations of a set of 1909 cryo-EM structures and quantified the agreement between the predicted and experimental cryo-EM data. For comparison, we also calculated density maps from the all-atom structures using the atomistic forward model. Comparing the *CC*_*mask*_ with the experimental maps given by the coarse-grained and atomistic forward models as a function of experimental resolution revealed that they perform equally well at lower resolutions (>4.0 Å), while the atomistic forward model is more accurate at higher resolutions (<4.0 Å) (Fig. 3). These results confirm the accuracy of our coarse-grained forward model and suggest that Martini models may be useful for single-structure refinement of medium-low resolution cryo-EM and cryo-ET data. Additionally, our new forward model will be useful for validating and biasing Martini simulations using cryo-EM and cryo-ET data. However, it should be noted that it is unlikely that Martini coarse-grained models could be directly used for model refinement without modification to the elastic network that Martini uses to limit the flexibility of such models. The use of coarse-grained models during single-structure and ensemble refinement may therefore require informed modifications to the elastic network or the use of other structure-based models (i.e., Go-models) to allow flexibility during sampling.

**Fig. 3.**
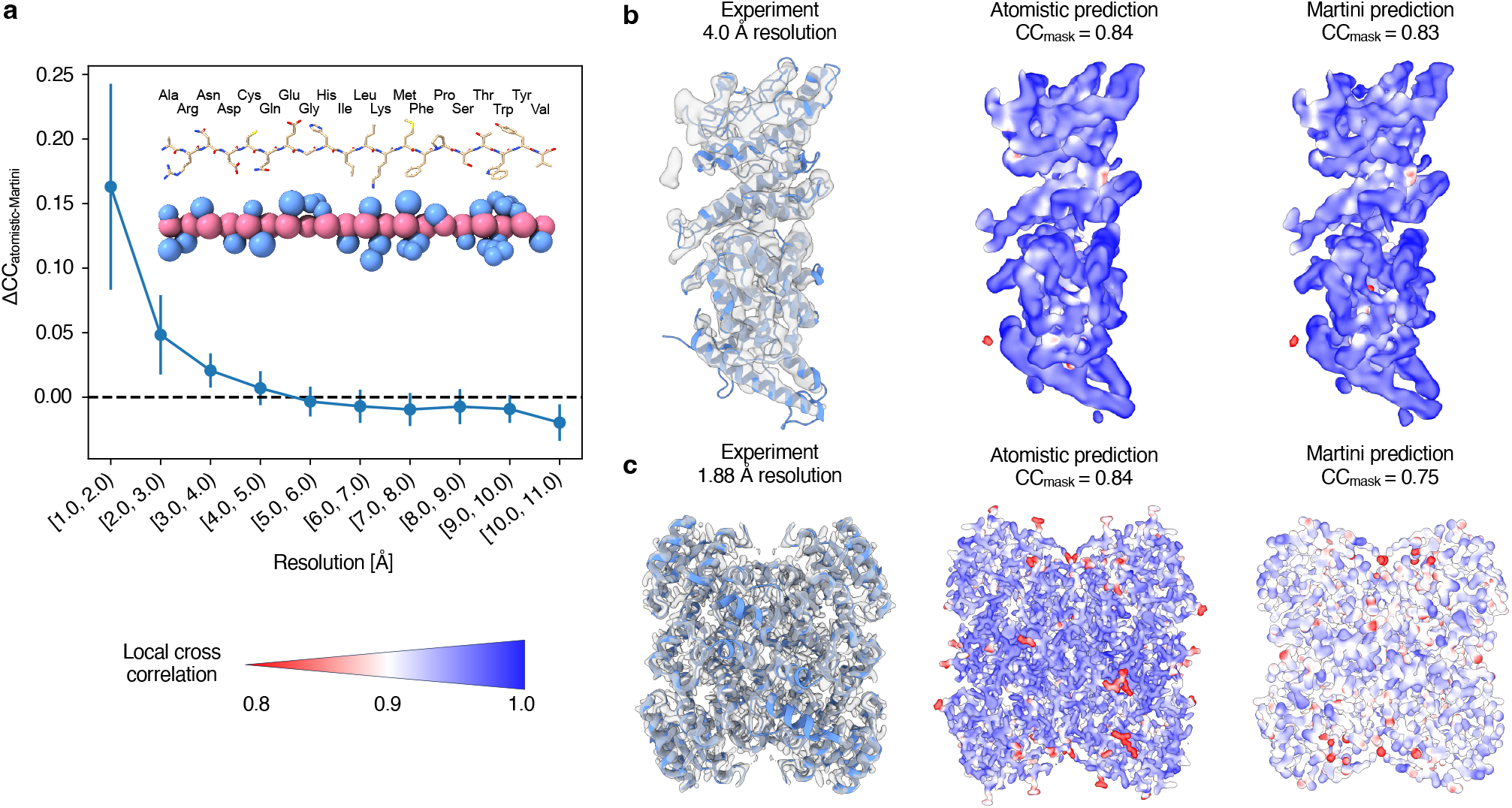
EMMIVox coarse-grained refinement. **a)** Difference between atomistic and coarse-grained (Martini 3) forward model in the agreement with 1909 experimental cryo-EM density maps. The plot shows the average difference in *CC*_*mask*_ (Δ*CC*) given by the two forward models as a function of the experimental resolution. Error bars show the standard deviation. **b-c)** Example structures and cryo-EM density maps at medium (**b**: PDB 7nnh^94^, 4.0 Å) and high (**c**: PDB 7o6q^95^, 1.88 Å) resolution. From left to right: experimental density maps and density maps predicted by the atomistic and coarse-grained forward models. The predicted maps are colored by local cross correlation to the experimental map (low: red, high: blue) and the global *CC*_*mask*_ with the experimental map is shown.

### Ensemble refinement

When coupled with metainference^41^, EMMIVox can be used to model structural ensembles by interpreting low resolution areas of a cryo-EM map in terms of a mixture of conformational heterogeneity and noise (Materials and Methods). To determine the accuracy of this combined approach, we determined structural ensembles for all the systems in our single-structure refinement benchmark set and compared their quality of fit to the experimental map to the EMMIVox single-structure model (Fig. 4A and S10). The *CC*_*mask*_ scores for structural ensembles had a median increase of 13% compared to single-structure models, reaching up to a *CC*_*mask*_ of 0.95 for the ensemble of Escherichia coli PBP1b (3.28 Å, PDB 7lq6). The largest improvement was observed for the Bacteriophage SPP1 (4 Å, PDB 6yeg), whose *CC*_*mask*_ score increased by 28.4% from 0.60 observed in the single-structure model to 0.77 for the ensemble.

**Fig. 4.**
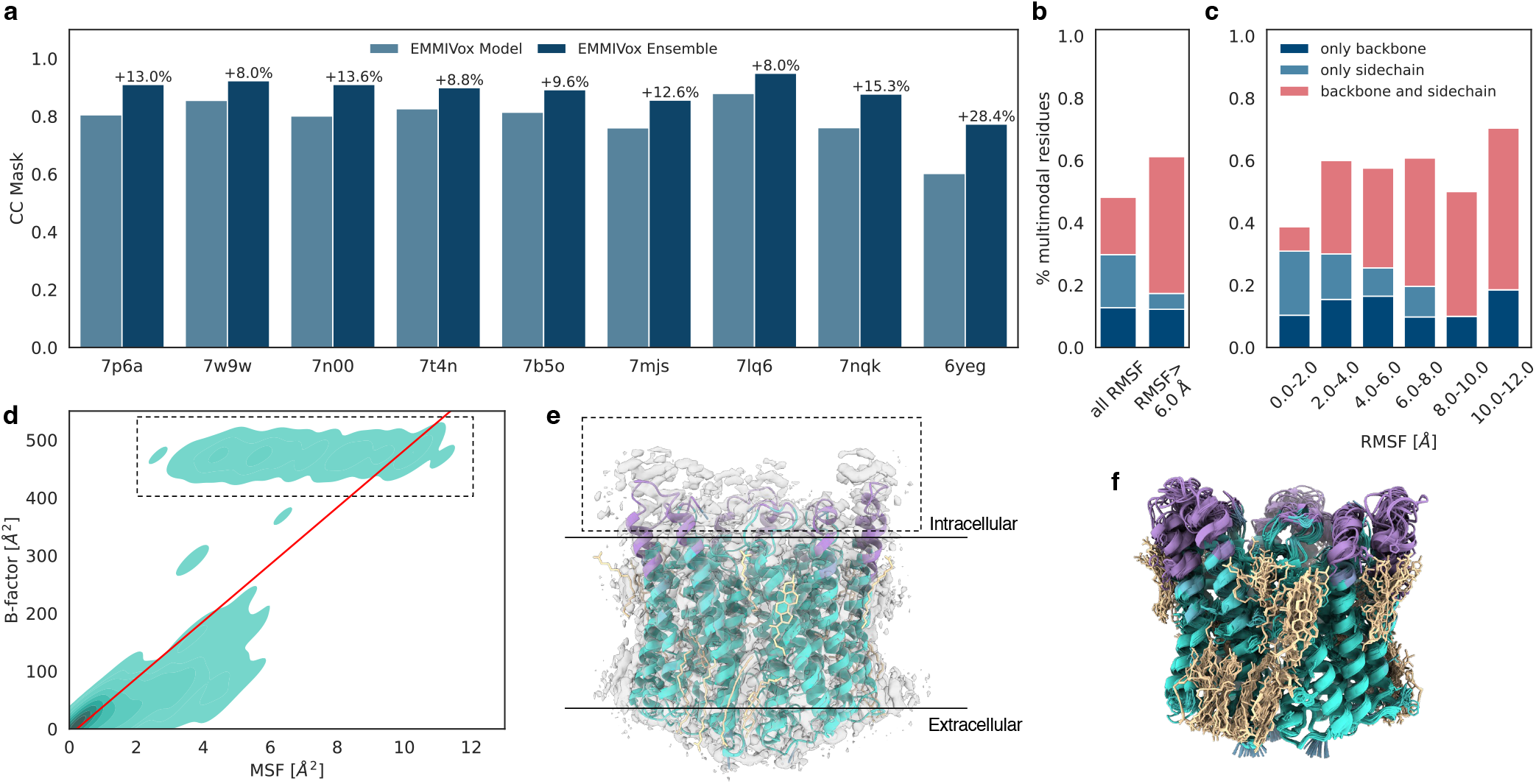
EMMIVox ensemble refinement benchmark. **a)** Model fit to experimental data (*CC*_*mask*_) of the nine systems in our benchmark set for the EMMIVox single-structure models (light blue) and EMMIVox ensemble (dark blue). The percentage increase in ensemble *CC*_*mask*_ is reported on top of the dark blue bars. **b)** Percentage of residues across the entire benchmark set that in the EMMIVox ensembles display multimodal conformational distributions according to the Folding Test of Multimodality. Each bar is decomposed into percentage of residues with only backbone (dark blue), only sidechain (light blue), and both backbone and sidechain (light red) multimodal distributions. Percentages calculated on all residues (left bar) and only on residues displaying RMSFs greater than 6 Å (right bar) are displayed separately. **c)** Percentage of residues that display multimodal conformational distributions as a function of residue RMSF. Colors as in panel **b**). **d**) Relation between per residue B-factor in the single-structure EMMIVox model and residue MSF within the EMMIVox ensemble in the case of ChRmine channelrhodopsin (2 Å, PDB 7w9w). The red line indicates a Bayesian linear fit between these two quantities. The same analysis for all the other systems of our benchmark set is reported in Fig. S11. **e**) Cryo-EM density map of ChRmine channelrhodopsin (EMD-32377) overlaid to the EMMIVox single-structure model. Residues with large B-factors highlighted in the dashed box of panel **d**) are colored in violet. **f**) EMMIVox structural ensemble of ChRmine channelrhodopsin. Colors as in panel **e**).

The superior model-data fit is not necessarily due only to the larger number of parameters used in ensemble refinement, *i*.*e*. multiple conformations, because single-structure models can implicitly account for dynamic effects using residue-level B-factors to describe fluctuations around an average conformation. The improved ensemble *CC*_*mask*_ suggests therefore that the dynamics of flexible regions might not be well represented by such Gaussian fluctuations. To investigate this point further, we examined in more detail the nature of the structural diversity observed in our EMMIVox ensembles. Across our entire benchmark set, 48% of residues showed a dynamic behavior that cannot be described by a unimodal distribution (Fig. 4B), neither at the backbone nor side-chain level. When restricting our analysis to medium-to-large scale dynamics, defined by residue Root Mean Square Fluctuations (RMSF) greater than 6 Å, the majority of residues (61%) sampled multimodal conformational distributions impacting a larger portion of backbone atoms as dynamics increases (Fig. 4C). These results, combined with the higher ensemble *CC*_*mask*_, indicate that unimodal, Gaussian-like distributions centered around a single-structure model with fluctuations proportional to the B-factors cannot accurately describe the conformational heterogeneity of the backbone and side chains that are averaged out in cryo-EM maps.

To determine whether the EMMIVox ensembles overinterpret experimental noise as conformational heterogeneity, we compared for each system the per-residue B-factors of the single-structure model to the residue MSF within the EMMIVox ensemble (Fig. S11). For all systems we observed a correlation between these two quantities indicating that the majority of the (multi-modal) heterogeneity observed in our ensembles corresponds to an increased B-factor in the single-structure model. However, most systems contained several residues with large B-factors that were not matched by proportionately large MSFs. This suggests that fuzzy densities around these residues correspond to experimental noise rather than structural dynamics. One intriguing example is ChRmine channelrhodopsin (2 Å, PDB 7w9w). In the EMMIVox single-structure model we observed disproportionately high B-factors for residues 191-214 and 269-279 (Fig. 4D, dashed box), which are located on the intracellular side (Fig. 4E). Density around these residues is extremely fuzzy, most likely due to the residual density from the antibody introduced to facilitate image alignment and/or radiation damage as this region is unprotected by the micelle during data collection (Fig. 4E). EMMIVox correctly did not overinterpret this fuzzy density as an extremely dynamic region but generated an ensemble with reduced conformational heterogeneity compared to the single-structure B-factors (Fig. 4F). This analysis, along with the previous observation that structural variety is often described by multimodal distributions, show that it is possible to get additional information about conformational heterogeneity beyond simply representing the continuous dynamics averaged in cryo-EM maps as Gaussian-like fluctuations proportional to the B-factors.

### Structural dynamics from atomistic cryo-EM maps: the GPT type 1a tau filament

As cryo-EM approaches atomistic resolution the question arises as to whether ensemble descriptions of density maps are still necessary^54^. We argue that ensemble representations are essential to extract all the information contained in these maps, such as the presence and location of semi-ordered water and lipids around proteins or local side chain flexibility, which cannot be captured by single-structure models nor by 2D particle classification. We demonstrate this concept using as example the GPT type 1a tau filament, which was recently determined at 1.9 Å resolution (PDB 7p6a, EMDB 13223)^40^.

Tau 1a proteins, which are thought to play a role in various neurodegenerative diseases, organize into tens-of-nm-long filaments with fold-dependent repeating structures. With EMMIVox, we determined the structural ensemble of the tau 1a filament from an atomistic cryo-EM map and observed several interesting features. First, the location of ordered water molecules found in the deposited PDB was correctly identified by the water density within the EMMIVox ensemble without the use of symmetry constraints (Fig. 5AB). Notably, we also observed additional water density close to the ordered waters, which was not present in the deposited PDB. This density corresponds to a second hydration shell composed of semi-ordered waters in exchange with bulk solution. Second, it is known that the aromatic residues at the surfaces of fibrils may be important to fibril growth^55^. Recent NMR studies revealed that the dynamics of these aromatic rings vary significantly depending on the amino acid location, with surface exposed amino acids populating a larger number of conformations in rapid exchange compared to aromatics within the fibril core^56^. This behavior was also observed in our EMMIVox ensemble (Fig. 5CD). This raises the question as to whether other non-aromatic surface exposed amino acids occupy multiple conformations, and if this information can be extracted from atomistic cryo-EM maps. We calculated the population of the conformers of two surface-exposed residues from the EMMIVox ensemble: K343, which in the deposited PDB was modeled in two alternative conformations with equal occupancy, and K347. We confirm that K343 occupies two distinct conformations with equal probability of 50% (Fig. 5E). Interestingly, we also observed a previously unmodeled minor conformation for K347 with an occupancy of 30% compared to 70% for the major conformation present in the deposited PDB (Fig. 5F).

**Fig. 5.**
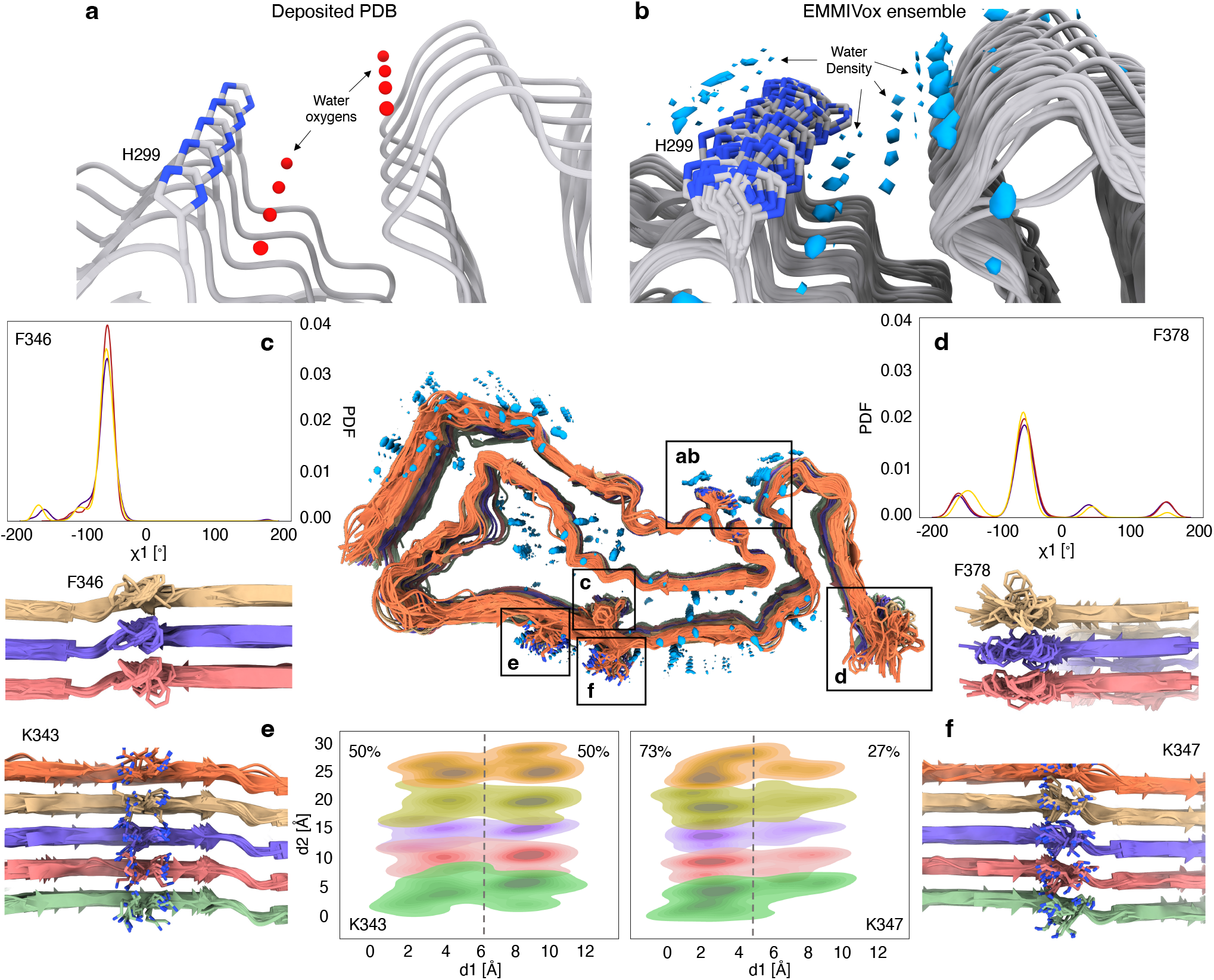
Case study: the GPT type 1a tau filament. **a)** Ordered waters in proximity of H299 resolved in the deposited structure (PDB 7p6a, 1.9 Å resolution, EMDB 13223). **b**) Water density within the EMMIVox ensemble showing both ordered and semi-ordered molecules in the proximity of H299. **cd**) Distribution of the χ1 sidechain dihedral angles of residues F346 (**c**) and F378 (**d**) computed from the EMMIVox ensembles, along with representative conformations. These distributions indicate that exposed aromatic residues populate a larger number of conformations compared to buried aromatics, as previously observed in NMR studies of other fibrils. **ef**) Structural ensembles of residues K343 (**e**) and K347 (**f**) indicating the presence of multiple sidechain conformations. In the EMMIVox ensemble, K343 occupies two distinct states with equal populations, as illustrated by the distribution of the lysine sidechain nitrogen positions projected on a plane parallel to the fibril surface. Both states were modelled in the deposited PDB with equal occupancy. K347 instead was modelled in a single conformation in PDB 7p6a, while in the EMMIVox ensemble it occupies a second minor conformation, populated only by 27%.

## Discussion

Here we presented EMMIVox, a tool to determine accurate single-structure models as well as structural ensembles by combining MD simulations driven by state-of-the-art molecular mechanics force fields and cryo-EM density maps. EMMIVox incorporates cryo-EM data in the form of voxels using a Bayesian framework that explicitly accounts for data correlation and noise. The method automatically balances model fit to experimental data with stereochemical quality and produces single-structure models that outperform the structures deposited in the PDB in terms of multiple key metrics. EMMIVox systematically reduced the number of serious clashes observed in deposited structures and optimized the positions of backbone and side-chain atoms to improve the description of physico-chemical interactions fundamental for protein stability, such as hydrogen bonds and salt bridges. These single-structure models can be leveraged for all those applications that require high-quality data, such as *in silico* structure-based drug design or training machine learning approaches.

Even more importantly, EMMIVox can be used to determine structural ensembles that reveal the conformational heterogeneity hidden in low-resolution areas of cryo-EM maps. While 2D classification, manifold embedding and other advanced image processing techniques^57-64^ have the potential to identify distinct conformational states directly from the single-particle data, we have shown here that even atomistic maps still present regions in which dynamics of flexible regions is averaged out. EMMIVox relies on both the cryo-EM data and physico-chemical knowledge to determine structural ensembles that capture key information that is lost in single-structure models, such as the presence and population of minor sidechain conformational states, semi-ordered waters, lipids and ligands. Our analysis indicated that this structural variability often corresponds to multimodal conformational distributions that cannot be well described by Gaussian-like fluctuations around an average model and proportional to single-structure B-factors. While these might appear as minor details, we have shown in the past that dynamics of flexible regions extracted from cryo-EM maps often play a crucial role in biological function, for example for ligand recognition in the ASCT2 transporter^65^ and to understand the effect of post-translational modifications in microtubules^66^.

EMMIVox presents multiple advantages with respect to our previous approach for structure determination from cryo-EM maps (EMMI)^35^. EMMIVox directly utilizes the density voxels of 3D cryo-EM density maps, replacing the previous representation method using Gaussian Mixture Models (GMM). While representing the cryo-EM density as voxels requires careful removal of correlated datapoints, it significantly increases usability, especially with atomistic cryo-EM density maps, for which determining accurate GMMs is prohibitively expensive. The direct use of voxels also enables estimation of experimental errors from deposited half-maps and the use of noise models for each density voxel, ensuring that various sources of error are not interpreted as structural dynamics when modeling structural ensembles. Additionally, our novel Bayesian inference approach to determine B-factors, which is unique to EMMIVox, enables accurate single-structure refinement and quantitative comparison of the models to the deposited PDBs using standard metrics of quality.

In spite of significant improvements, EMMIVox still presents some limitations. First, the computational cost of EMMIVox is higher compared to the real-space refinement approach implemented in PHENIX. The main reason is the use of accurate models of the environment, *i*.*e*. explicit water molecules, lipid bilayer, and solution ions, and of the intermolecular forces, especially long-range interactions. Furthermore, while single-structure refinement can be performed on a GPU-enabled Desktop computer in 10 to 24 hours, the determination of structural ensembles requires simultaneous use of multiple computer nodes, which may be out of reach for some users. Second, EMMIVox was designed to capture the underlying conformational distribution of the selected single-particle images used to reconstruct the 3D density map. As this data is a subset of the entire particle set, it is important to note that the resulting ensemble may represent local dynamics around a stable macrostate and not the entire conformational landscape. Furthermore, it should be kept in mind that the cryo-EM cooling process might perturb the room temperature ensemble by reducing thermal motion and enabling equilibration into lower free-energy conformations^67^. In EMMIVox, these issues are partially mitigated by using the room temperature ensemble provided by the molecular mechanics force field as a prior. Third, while EMMIVox has been shown to accurately and rapidly capture conformations present in available cryo-EM density maps, including those which are difficult to observe on short time scales in standard MD simulations (Fig. S12), EMMIVox may have trouble sampling conformations separated by large free-energy barriers. In this case, sampling issues can be alleviated by combining EMMIVox with various enhanced sampling techniques, such as metadynamics as recently proposed in the MEMMI approach^68^. Finally, because of these sampling limitations, an initial model that already fits the experimental density to a good extent is required as starting point. AlphaFold2^9^, ab initio machine learning and other automated model building techniques^31, 69^, or flexible fitting methods such as MDFF^21^, TEMPy-ReFF^22^, and the novel maximum likelihood approach implemented in GROMACS^24^, can provide excellent starting models for accurate single-structure and/or ensemble refinement with EMMIVox.

Despite these limitations, EMMIVox enables determining accurate structural and dynamic models using *in vitro* data, and holds promise as a valuable tool for future efforts focused on structure determination *in situ*. As the resolution of cryo-ET improves (Fig. S1B), high-quality in cell data will become readily available and integrative approaches that combine various sources of experimental and computational data will be required to obtain accurate structural models. The coarse-grained forward model implemented in EMMIVox will enable refining models of large macromolecular architectures from sub-nanometer subtomogram averaging data. Additionally, the Bayesian framework on which EMMIVox is built enables automatic weighting of multiple sources of *in silico* and experimental data, making it a perfect framework for integrative structure and dynamic determination in cell. This integration of a diverse set of data is further facilitated by the implementation of EMMIVox in the PLUMED-ISDB^70^ module of PLUMED^38^, which makes it possible to combine a wide range of different types of experimental data.

In summary, EMMIVox represents a flexible instrument to convert cryo-EM data into high-quality structural models. Our approach can be used to reveal dynamic properties of proteins, lipids, ligands, waters, and ions by extracting information from cryo-EM density maps that would otherwise be lost. These models can advance our understanding of the molecular mechanisms that drive biological functions, and provide valuable information for structure-based drug design and training of novel machine learning approaches. Looking ahead, we envision that EMMIVox will be an indispensable part of integrative structural biology pipelines and will contribute to obtaining a more complete picture of highly intricate and dynamic systems in biologically relevant environments.

## Materials and Methods

### Theory of EMMIVox for single-structure refinement

#### General overview

EMMIVox is based on a Bayesian inference framework^37^ that estimates the probability of a model *M* given the information available about the system, including prior physico-chemical knowledge and newly acquired experimental data. The posterior probability *p*(*M*|*D*) of model *M*, which is defined in terms of its structure *X* and other parameters, given data *D* and prior knowledge is:

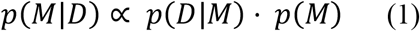

where the *likelihood function p*(*D*|*M*) is the probability of observing data *D* given *M* and the *prior p*(*M*) is the probability of model *M* given the prior information. At variance with the previous EMMI approach^29, 35^, here we define the experimental data as a set of density values *D =* {*d*_*i*_} observed in the voxels of a cryo-EM three-dimensional (3D) map. To define the likelihood function, one needs *i)* a *forward model f*_*i*_(*X*) to predict the density that would be observed in voxel *i* for structure *X* in absence of noise, and *ii)* a *noise model* that defines the distribution of deviations between observed and predicted data.

#### Atomistic forward model

To predict the density in voxel *i* of a cryo-EM map from an atomistic model, we used the fast Fourier transform of a 5-Gaussian fit of the electron scattering factors^71^:

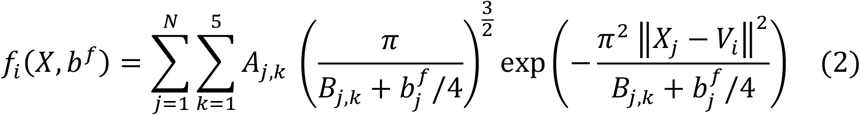

where *X* _*j*_ and *V*_*i*_ are the coordinates of atom *j* and the center of the voxel *i*, respectively. The Gaussians parameters *A*_*k*_ and *B*_*k*_ depend on the atom type (Tab. S1). The B-factors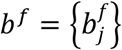, which are defined for each atom *j* but are identical within individual residues, enable smoothing the forward model prediction to describe low-resolution regions with fuzzy density using a single-structure model. The external sum runs over all the non-hydrogen atoms, with exclusion of the carboxylate oxygens of glutamic and aspartic acid, as these groups are often damaged by the electron beam. To reduce the computational cost, we further restricted this summation to a neighbor list of atoms with distance cutoff from the voxel center of 1.0 nm.

#### Noise model

We use a Gaussian noise model for the density *d*_*i*_ observed in voxel *i*.:

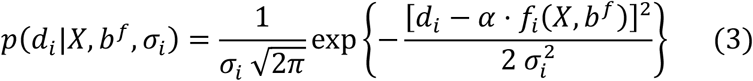

where the uncertainty parameter 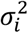 quantifies the level of experimental noise as well as errors in the forward model. These variable parameters allow to dynamically adjust the intensity of the cryo-EM spatial restraints during model refinement based on the accuracy in the data, which is inferred by the consistency between cryo-EM data, physico-chemical prior and potentially additional experimental data. High-noise voxels (*outliers*) are therefore automatically identified and downweighed in the refinement of the structural model in favor of the prior information. *α* is a constant scaling factor between the entire set of voxels of the predicted and experimental maps, and it is optimized prior to production runs (*Scale factor optimization*).

#### Data correlation and likelihood function

The data likelihood *p*(*D*|*M*) is typically expressed as a product of individual likelihoods *p*(*d*_*i*_|*M*), one for each experimental data point, under the assumption that these points are independent. In case of cryo-EM data, this assumption does not hold because neighboring voxels of a 3D map contain correlated information. Ignoring such correlation would lead to overcounting the number of independent data points and ultimately biasing the refinement towards overfitting the data at the expenses of the stereochemical quality of the models. To reduce data correlation, we developed the following pre-filtering procedure:

1. We first selected all the voxels of the 3D cryo-EM map that are within 3.5 Å of the atoms in the PDB structure. Density voxels with negative values were discarded. The selected atoms include protein atoms as well as ordered waters, ions, small-molecules and lipids. We sorted the selected voxels in descending order based on the value of the density: this constitutes our initial pool of data points *D =* {*d*_*i*_};
2. We started from the first voxel *d*_1_ (highest density) in pool *D* and calculated the density autocorrelation function for displacements up to a few voxels in the *x, y*, and *z* direction starting from this location. For each displacement, the autocorrelation function is calculated by averaging over multiple starting voxels in a cubic minibox of side equal to 4 Å and centered on *d*_1_. Equivalently, the autocorrelation can be expressed as the Pearson correlation coefficient of a series of density values and its space-lagged version;
3. We removed from pool *D* all the voxels within the cubic minibox with Pearson correlation coefficient greater than a predefined threshold, with the exception of *d*_1_.
4. We moved to the next voxel *d*_2_ in pool *D* in descending order of density and re-applied the filtering procedure of step 2 and 3;
5. We iterated until reaching the voxel in pool *D* with lowest density.

It should be noted that sub-sampling procedures are commonly used in structural modelling with experimental data, for example to remove correlation between points in Small-Angle X-ray scattering profiles^72^. PHENIX itself utilizes only the density in the positions occupied by the atoms during each step of refinement, which is obtained by interpolating the density in the voxels surrounding each atom^23^. After pre-filtering the map with the procedure describe above, we assume that the *N*_*v*_ selected voxels can be considered as independent data points and write the total likelihood function as:

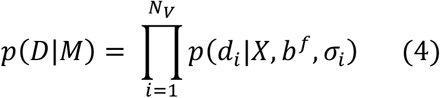

#### Priors

Experimental cryo-EM density maps may contain errors from various sources including systematic errors during 3D reconstruction, noise from the use of a limited number of images taken at various sample orientations, and radiation damage from exposure to the electron beam. Furthermore, the forward model used to predict a map from a structural model is intrinsically inaccurate. While these errors might be difficult to measure, it is critically important that they are not misinterpreted as structural dynamics during the generation of structural ensembles. To guide the inference of the uncertainty parameters *σ*_*i*_ that quantify the noise level in the map, we defined a lower bound 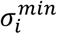 for each voxel by calculating the voxel-by-voxel variation between two independent reconstructions, or *half maps*. While this quantity does not describe all the sources of error at play, we expect that the total error in a given voxel cannot be lower than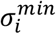 . We imposed this lower bound using the following prior for the uncertainty parameters *σ*_*i*_^73^:

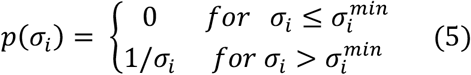

where 1/*σ*_*i*_ is a typical Jeffreys prior.

Inspired by PHENIX, we also added a restraint to guide B-factors inference and avoid that residues close in space have B-factors significantly different from each other. These restraints are enforced by:

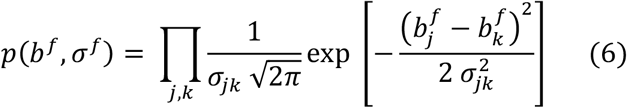

where the product is over all pairs of B-factors corresponding to residues for which at least a pair of atoms is closer than 5 Å. At variance with the B-factor restraint implemented in PHENIX, our Bayesian approach tolerates outliers, *i*.*e*. pair of close residues with significantly different B-factors, via the introduction of the uncertainty parameters *σ*^*f*^ *=* {*σ*_*jk*_}. This approach allows us to better model during single-structure refinement those regions of the system that undergo a sudden transition between order and disorder.

Finally, as structural prior *p*(*X*) we used state-of-the-art atomistic force fields *E*_*FF*_(*X*), which enabled us to accurately model the system as well as the environment, including explicit water molecules, ions, small-molecules, and lipids:

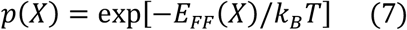

where *k*_*B*_ is the Boltzmann constant and *T* the temperature of the system.

#### Marginalization

To avoid sampling the uncertainty parameters *σ*_*i*_ and *σ*^*f*^, we marginalized the corresponding distributions. The resulting marginal data likelihood is:

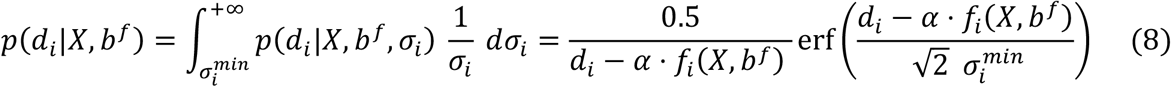

while the B-factors prior becomes upon introduction of a Jeffreys prior 1/*σ*_*jk*_:

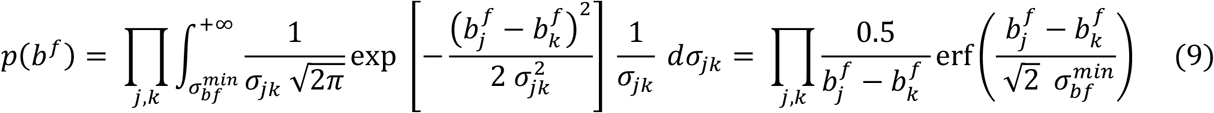

where 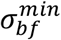 is a lower bound for the B-factor uncertainty parameters set equal to 0.1 nm^2^.

#### EMMIVox hybrid energy function

After defining all the component of our approach and marginalizing the uncertainty parameters, we obtain the final EMMIVox posterior distribution:

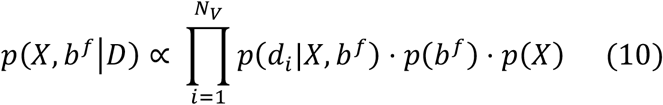

where the marginal data likelihood is given by Eq. 8, the B-factors prior by Eq. 9, and the structural prior by Eq. 7. To sample the posterior, we define the associated hybrid energy function as:

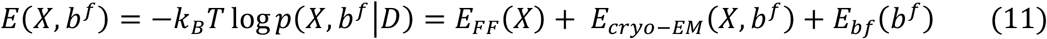

The EMMIVox hybrid energy is therefore decomposed into: *i)* the molecular mechanics force field *E*_*FF*_ (Eq. 7), *ii)* the spatial restraints *E*_*cryo-Em*_ to enforce the model agreement with the cryo-EM map (Eq. 8), and *iii)* the restraints *E*_*bf*_ to guide B-factors determination (Eq. 9). A Gibbs sampling scheme is used to sample model coordinates with Molecular Dynamics (MD) and B-factors parameters with Monte Carlo (MC).

### Development of the coarse-grained forward model

#### Parameterization

To enable the use of EMMIVox with coarse-grained representations, we developed a forward model to calculate density maps from Martini 3 protein structures^53^. We parameterized the model by fitting to predictions of the atomistic forward model. In the atomistic forward model, the atomic electron scattering factors are given by the Fourier transform of a 5-gaussian mixture (see section *Atomistic forward model* and Eq. 2). We used the same framework for the Martini forward model, but with bead electron scattering factors given by a single Gaussian, *i*.*e*. only two parameters *A*_*k*_ and *B*_*k*_ for each Martini bead type *k*. We determined an individual set of *A*_*k*_ and *B*_*k*_ for each bead position in each type of amino acid for a total of 52 different bead types using the following procedure. To capture the conformational variation in the atoms mapping to each bead type, we used a set of 2906 structures to fit the parameters. These were all the structures obtained with single-particle cryo-EM in the period 2020-2023, with resolution ranging from 1.2 Å to 11 Å. We divided the structures into groups of atoms, each corresponding to one Martini bead type, skipping any beads with missing atoms. For each group, we defined a cubic box of voxels centered on each atom and with side equal to 6 Å (for a total of 6859 evenly spaced voxels). In this set of voxels, we calculated a density map from the group of atoms using the atomistic forward model and from the corresponding Martini bead using the coarse-grained forward model with a grid scan of *A*_*k*_ and *B*_*k*_. We scanned values of *A*_*k*_ from 0 to 8.0 in steps of 0.1 and *B*_*k*_ from 0 Å^2^ to 40.0 Å^2^ in steps of 0.5 Å^2^. We evaluated the agreement between the two forward models as a function of *A*_*k*_ and *B*_*k*_ using the squared error summed over all the voxels:

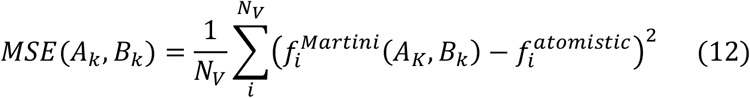

where *N*_*V*_ is the number of voxels, 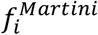 and 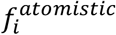 are the voxel densities predicted by the Martini and atomistic forward models, respectively. For each of the 52 Martini bead types, we averaged the MSE grid scan over the bead type’s occurrences in all 2906 structures and selected the set of *A*_*k*_ and *B*_*k*_ that minimized the average MSE (Tab. S2).

#### Validation

To validate the Martini forward model and compare its accuracy with that of the atomistic forward model, we evaluated the agreement between predicted and experimental density maps for 1909 cryo-EM structures. These were selected from our initial set of 2906 structures as those that could be easily mapped to the Martini 3 representation using the *martinize2* python script without any additional modifications to the structures^74^. We calculated density maps from the atomistic and Martini representations using the respective forward models and calculated the *CC*_*mask*_ with the experimental density maps. We used the same protocol described below (in *Details of the single-structure refinement benchmark*) to preprocess the experimental maps and to optimize the B-factors and scale factors of the predicted maps to maximize *CC*_*mask*_. The B-factors were sampled for 1000 MC steps and the scale factor was scanned from 0.7 to 1.3 in steps of 0.05. No data correlation filtering was used for the experimental maps. To evaluate the difference between the forward models as a function of the experimental resolution, we binned the structures by resolution (bin size equal to 1.0 Å) and calculated for each bin the average difference in *CC*_*mask*_ between the atomistic and Martini forward models:

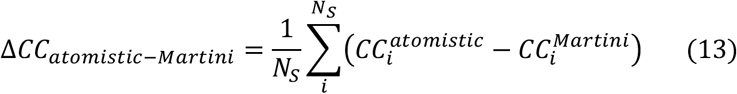

where *N*_*S*_ is the number of structures in the bin.

### Theory of EMMIVox for ensemble refinement

To model structural ensembles fitting a cryo-EM density map, we coupled EMMIVox with metainference^41^. Metainference is a general Bayesian inference approach that enables modelling structural ensembles using any kind of ensemble-averaged experimental data as well as prior physico-chemical information. Inspired by the maximum entropy/replica-averaged approach^75^, the metainference posterior is expressed in terms of several copies of the system (or *replicas*), which represent the conformational heterogeneity of the system. With a Gaussian data likelihood, as in the EMMIVox case, the metainference posterior is:

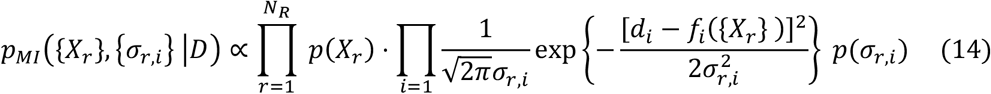

where {*X*_*r*_} are the structures of the *N*_*R*_ replicas of the system and {*σ*_*r,i*_} the error parameters, one per replica *r* and experimental data point *i*. The predicted experimental observable *f*_*i*_({*X*_*r*_}) is calculated as the average of *f*_*i*_ over the *N*_*R*_ replicas. Sampling of the metainference posterior is performed by a multi-replica MD/MC simulation driven by the hybrid energy function associated to Eq. 14. This approach has been extensively used to determine structural ensembles of highly dynamic and disordered systems using NMR spectroscopy^76, 77^ as well as SAXS/SANS^78, 79^ data. Metainference was also used to determine structural ensembles from cryo-EM data in our previous EMMI approach based on a Gaussian Mixture Models representation of the cryo-EM map^29, 35^. In combination with EMMIVox and the more accurate noise models developed here, metainference enables ensemble-refinement by interpreting low resolution areas of a cryo-EM map in terms of a mixture of conformational heterogeneity and noise.

### Details of the single-structure refinement benchmark

#### Details of the systems

Nine systems determined at a resolution ranging from 1.9 Å to 4.0 Å were chosen to benchmark the EMMIVox single-structure refinement (Tab. S3): the GPT type 1a tau filament, the ChRmine channelrhodopsin, the Anaplastic lymphoma kinase extracellular domain fragment in complex with an activating ligand, a mutant of dimeric unphosporylated Pediculus humanus protein kinase, the human CDK-activating kinase bound to the clinical inhibitor ICEC0942, the major facilitator superfamily domain containing 2A in complex with LPC-18:3, the Escherichia coli PBP1b, the mammalian peptide transporter PepT2, and the SPP1 bacteriophage tail tube.

#### Setup and general MD details

Missing residues in the deposited PDB were modelled using GalaxyFill^80^ or, in case missing residues could not be placed, with Modeller^15^ v. 10.1. The resulting model was then processed using the CHARMM-GUI^81^ server. Membrane proteins (PDB ids 7w9w, 7mjs, and 7nqk) were inserted in a homogeneous POPC lipid bilayer. Each system was solvated in a triclinic box with dimensions chosen in such a way that the edge of the box was 1.0 nm away from the closest model atom. K+ and Cl-were added to ensure charge neutrality at concentration equal to 0.15 M. The CHARMM36m^82^ forcefield was used for proteins and lipids, CgenFF for small molecules^83^, and the mTIP3P^84^ model for water molecules. In all simulations the equations of motion were integrated by a leap-frog algorithm with timestep equal to 2 fs. The smooth particle mesh Ewald^85^ method was used to calculate electrostatic interactions with a cutoff equal to 1.2 nm. Van der Waals interactions were gradually switched off at 1.0 nm and cut off at 1.2 nm. All simulations were carried out using GROMACS^86^ v. 2020.5 equipped with the development version of PLUMED^38^. To optimize performances, EMMIVox is implemented in PLUMED with libtorch^87^ to efficiently calculate the cryo-EM forward model and the hybrid energy function on the GPU.

#### Cryo-EM map preprocessing

For each system, we downloaded the cryo-EM map as well as the two half maps from the EMDB database. We applied our pre-processing procedure to select the voxels within 3.5 Å of the model atoms and filter them to reduce data correlation, with a threshold of 0.7, 0.8, 0.9, and 1.0 (no filtering). The two experimental half maps were used to calculate the lower bound for the density uncertainty parameters. Since CHARMM-GUI processing of the deposited PDB often translates and rotates the input conformation, we calculated the transformation that aligns initial and final models and applied it to all the voxels selected for refinement. Finally, a single map data file was created with the list of voxels to be used by PLUMED for refinement and, for each voxel, the following information: (transformed) coordinates of the voxel center, density value, and uncertainty lower bound.

#### Equilibration

Energy minimization was performed on each system using the steepest decent approach. A 1 ns-long NPT equilibration was then performed with the Bussi-Donadio-Parrinello thermostat^88^ and the Berendsen barostat^89^, set at 300 K and 1 atm respectively. In systems containing a lipid bilayer, the pressure coupling type was semi-isotropic to allow deformations in the *xy* plane independent from the *z*-axis. Next, NVT simulations were carried out for 2 ns using the Bussi-Donadio-Parrinello thermostat at 300 K. During the last two steps of equilibration, positional restraints were applied to all the heavy atoms of the protein and any other components contributing to the predicted cryo-EM density map.

#### Modelling of ordered waters

In presence of ordered waters in the deposited PDB, a special treatment was required. First, resolved waters as well as all the water molecules within 3.5 Å in the initial CHARMM-GUI solvated model (*buffer waters*) contributed to the map prediction via our forward model. Second, the positions of ordered waters were restrained during equilibration; the buffer waters were restrained to stay within 8 Å from a reference protein atom, defined as the closest to each water molecule in the initial model.

#### Scale factor optimization

To reduce the number of free parameters to sample during the production run, we determined an optimal scaling factor between predicted and observed cryo-EM maps. We analyzed with the PLUMED *driver* tool the trajectory obtained with positional restraints during the NVT equilibration for different values of the scaling factors in the range from 0.5 to 1.5 at intervals of 0.05. For each value of the scaling factor, we sampled with MC the B-factors while re-reading the trajectory, with a maximum MC move per B-factor equal to 0.05 nm^2^. To optimize sampling, B-factors were initialized using an empirical relationship between map resolution (*res*, in nm) and average B-factor values (in nm^2^) determined from 8000 deposited PDBs obtained from cryo-EM maps with resolution less than 5 Å:

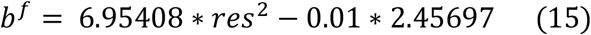

The scaling factor that resulted in the lowest EMMIVox hybrid energy along the entire trajectory was selected to be used in the production simulations described in the following section.

#### Production

All single-structure refinement production simulations were performed in the NVT ensemble with Bussi-Donadio-Parrinello thermostat at 300 K for 20 ns (NPT ensemble with Parrinello-Rahman barostat^90^ at 1 atm for membrane proteins). Before calculating the EMMIVox hybrid energy, structural discontinuities due to Periodic Boundary Conditions (PBC) were fixed on-the-fly by PLUMED. Furthermore, to optimize performances, we: *i)* updated the forward model neighbor list every 50 MD steps; *ii)* sampled the B-factors every 500 MD steps with maximum MC move equal to equal to 0.05 nm^2^; *iii)* used a multiple-time step algorithm for PBC reconstruction and EMMIVox hybrid energy calculation with stride equal to 4 MD steps^91^. The system trajectory was saved to file every 10 ps for subsequent analysis. Ordered and buffer waters that contributed to the map prediction were restrained to stay within 8 Å from a reference protein atom, defined as the closest to each water molecule in the initial model. These restraints allowed: *i)* exchanges between the ordered molecules resolved in the PDB and their surrounding buffer waters; *ii)* to identify additional sites of (semi)ordered molecules in the proximity of the resolved waters, for example a second coordination shell.

#### Analysis

To generate the final single-structure refined model, we first extracted the conformation with lowest EMMIVox hybrid energy from the production run as well as the associated B-factors. We then performed an (EMMIVox hybrid) energy minimization using the steepest decent approach. During minimization, B-factors were sampled starting from those found in the initial model using a modified MC sampler in which only downhill moves were accepted. B-factors were sampled every 100 steps with a maximum MC move equal to 0.1 nm^2^. The forward model neighbor list was updated at every step. Next, we selected the conformation at the end of the minimization and fixed discontinuities in the structure due to PBC with the PLUMED *driver* tool. We then generated a PDB file containing only the heavy atoms of the systems that were used to predict the cryo-EM density map during production. This model was re-aligned to the original cryo-EM map downloaded from the EMDB using the inverse transformation computed in the section *Cryo-EM map preprocessing*. Finally, we added to the PDB the B-factors obtained at the end of the final minimization. MolProbity was used to compute the clashscore and MolProbity score, while PHENIX v. 1.15.2 was used to evaluate the fit to the experimental map with the EMRinger score. We implemented the calculation of *CC*_*mask*_^51^ in a GPU-enabled python script and generalized it to compute this score from a structural ensemble. At variance with PHENIX, we did not optimize an isotropic B-factor before the *CC*_*mask*_ calculation. Despite this difference, the *CC*_*mask*_ calculated by PHENIX and by our python script are strongly correlated (Fig. S13). For all the fit-to-data calculations, the original map as downloaded from the EMDB was used, regardless of the data correlation cutoff and the total number of voxels used in the refinement. For the analysis of physico-chemical fingerprints (Fig. 2), we used MDAnalysis^92^ v. 2.0 to calculate the number of hydrogen bonds from a single-structure model with donor-acceptor distance and angle cutoff equal to 3.0 Å and 150 degrees, respectively. Hydrogen bonds between anionic carboxylate (RCOO^−^) of either aspartic acid or glutamic acid and the cationic ammonium (RNH_3_^+^) from lysine or arginine were classified as salt bridges. Clashes between atoms with overlap between van der Waals radii greater than 0.4 Å were calculated with PHENIX.clashscore.

### Details of the ensemble refinement benchmark

#### Setup

We determined EMMIVox structural ensembles fitting the cryo-EM map for all the systems selected for the single-structure refinement benchmark. To prepare the ensemble simulations, we first extracted the conformation with lowest EMMIVox hybrid energy from the single-structure refinement production run and identified the minimum B-factor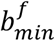 across all residues of this conformation. We then extracted 16 frames from the second half of the single-structure refinement production runs equally distributed in time. These conformations will be used as starting points of the production run described in the following section.

#### Production

EMMIVox ensemble simulations were performed with the same settings used in single-structure refinement, except for the fact that during ensemble refinement B-factors were not sampled but kept constant and all equal to the 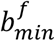 identified in the previous step. This value of B-factors can be considered as either a baseline dynamic of the most rigid residue of the system or a minimum noise level of the best resolved residue. In both cases, using such constant B-factor during ensemble refinement contributes to avoiding overinterpreting noise in terms of conformational heterogeneity. 16 metainference replicas were used, each one simulated for 20 ns, resulting in 320 ns of ensemble trajectory for each system.

#### Analysis

After the EMMIVox ensemble simulations were completed, the trajectories of all the metainference replicas were concatenated, resulting in the complete structural ensemble of the system. After fixing with PLUMED the structural discontinuities due to PBC, we re-aligned the EMMIVox ensemble to the original cryo-EM map downloaded from the EMDB using the inverse transformation computed in the section *Cryo-EM map preprocessing*. We then evaluated the fit to the data by calculating the *CC*_*mask*_ of the average cryo-EM map from the EMMIVox ensemble and the experimental map. To make a fair comparison, we selected the voxels for the *CC*_*mask*_ calculation based only on the single-structure model using the standard convention^51^. In this way, the calculation of *CC*_*mask*_ for single-structure and ensemble models was performed on the same set of voxels. The analysis of multimodality of the conformational distribution of each individual residue (Fig. 4) was performed using the Folding Test of Unimodality implemented in libfolding^93^ (https://github.com/asiffer/python3-libfolding).

## Software and data availability

EMMIVox is implemented as a part of the PLUMED-ISDB^70^ module in the development version (GitHub master branch) of PLUMED^38^ (https://github.com/plumed/plumed.github.io) and will be available in the official release starting from v. 2.10. The GROMACS topologies and PLUMED input files used in our benchmark as well as the EMMIVox refined single-structure models are available in PLUMED-NEST (www.plumed-nest.org), the public repository of the PLUMED consortium^39^, as plumID:23.041. Scripts to prepare and analyze EMMIVox simulations as well as complete tutorials for single-structure and ensemble refinement are available on GitHub (https://github.com/COSBlab/EMMIVox).

## Acknowledgements

M.B. and S.E.H. acknowledge the support of the French Agence Nationale de la Recherche (ANR), under grant ANR-20-CE45-0002 (project EMMI). S.E.H. is founded by a Roux-Cantarini fellowship from the Institut Pasteur (Paris, France). M.B. would like to acknowledge PRACE for awarding access to Piz Daint at CSCS, Switzerland.

## Ethics declarations

The authors declare no conflict of interests.

