## Supplementary Information for "Accurate model and ensemble refinement using cryo-electron microscopy maps and Bayesian inference"

### List of Supplementary Tables

| Atom type | $A_1$ | $A_2$ | $A_3$ | $A_4$ | $A_5$ | $B_1$ | $B_2$ | $B_3$ | $B_4$ | $B_5$ |
|---|---|---|---|---|---|---|---|---|---|---|
| C | 0.0489 | 0.2091 | 0.7537 | 1.142 | 0.3555 | 0.114 | 1.0825 | 5.4281 | 17.8811 | 51.134 |
| O | 0.0365 | 0.1729 | 0.5805 | 0.8814 | 0.3121 | 0.0652 | 0.6184 | 2.9449 | 9.6298 | 28.2194 |
| N | 0.0267 | 0.1328 | 0.5301 | 1.102 | 0.4215 | 0.0541 | 0.5165 | 2.8207 | 10.6297 | 34.3764 |
| S | 0.0915 | 0.4312 | 1.0847 | 2.4671 | 1.0852 | 0.0838 | 0.7788 | 4.3462 | 15.5846 | 44.63655 |
| P | 0.1005 | 0.4615 | 1.0663 | 2.5854 | 1.2725 | 0.0977 | 0.9084 | 4.9654 | 18.5471 | 54.3648 |
| F | 0.0382 | 0.1822 | 0.5972 | 0.7707 | 0.213 | 0.0613 | 0.5753 | 2.6858 | 8.8214 | 25.6668 |
| Na | 0.126 | 0.6442 | 0.8893 | 1.8197 | 1.2988 | 0.1684 | 1.715 | 8.8386 | 50.8265 | 147.2073 |
| Mg | 0.113 | 0.5575 | 0.9046 | 2.158 | 1.4735 | 0.1356 | 1.3579 | 6.9255 | 32.3165 | 92.1138 |
| Cl | 0.0799 | 0.3891 | 1.0037 | 2.3332 | 1.0507 | 0.0694 | 0.6443 | 3.5351 | 12.5058 | 35.8633 |
| Ca | 0.2355 | 0.9916 | 2.3959 | 3.7252 | 2.5647 | 0.1742 | 1.8329 | 8.8407 | 47.4583 | 134.9613 |
| K | 0.2149 | 0.8703 | 2.4999 | 2.3591 | 3.0318 | 0.166 | 1.6906 | 8.7447 | 46.7825 | 165.6923 |
| Zn | 0.178 | 0.8096 | 1.6744 | 1.9499 | 1.4495 | 0.0876 | 0.865 | 3.8612 | 18.8726 | 64.7016 |

**Tab. S1. Parameters of the forward model to predict a density map from an atomistic model.** These parameters are obtained from a 5-Gaussians fit of the electron atomic scattering factors for $s$ up to 6.0 Å$^{-1}$. $B_k$ parameters are expressed in Å$^2$.

| Bead type | $A_1$ | $B_1$ | Bead type | $A_1$ | $B_1$ | Bead type | $A_1$ | $B_1$ | Bead type | $A_1$ | $B_1$ |
|---|---|---|---|---|---|---|---|---|---|---|---|
| ALA_BB | 9.00 | 22.00 | HIS_BB | 9.50 | 23.00 | ASN_BB | 9.00 | 22.00 | VAL_BB | 9.50 | 23.00 |
| ALA_SC1 | 0.50 | 0.50 | HIS_SC1 | 4.50 | 11.50 | ASN_SC1 | 9.00 | 18.50 | VAL_SC1 | 7.00 | 18.00 |
| CYS_BB | 9.50 | 23.00 | HIS_SC2 | 4.00 | 9.00 | PRO_BB | 9.50 | 23.50 | TRP_BB | 9.50 | 23.00 |
| CYS_SC1 | 5.50 | 8.50 | HIS_SC3 | 4.00 | 8.50 | PRO_SC1 | 7.00 | 17.50 | TRP_SC1 | 4.50 | 11.50 |
| ASP_BB | 9.50 | 23.00 | ILE_BB | 9.50 | 23.00 | GLN_BB | 9.00 | 22.00 | TRP_SC2 | 4.00 | 9.00 |
| ASP_SC1 | 8.50 | 17.00 | ILE_SC1 | 10.00 | 25.50 | GLN_SC1 | 11.50 | 24.50 | TRP_SC3 | 4.50 | 11.00 |
| GLU_BB | 9.00 | 22.00 | LYS_BB | 9.00 | 22.00 | ARG_BB | 9.50 | 23.00 | TRP_SC4 | 4.50 | 11.00 |
| GLU_SC1 | 11.50 | 24.00 | LYS_SC1 | 7.00 | 18.00 | ARG_SC1 | 7.00 | 18.00 | TRP_SC5 | 4.00 | 9.50 |
| PHE_BB | 9.50 | 23.00 | LYS_SC2 | 4.50 | 11.00 | ARG_SC2 | 9.00 | 18.00 | TYR_BB | 9.50 | 23.00 |
| PHE_SC1 | 7.00 | 17.50 | LEU_BB | 9.00 | 22.00 | SER_BB | 9.50 | 23.00 | TYR_SC1 | 4.50 | 12.00 |
| PHE_SC2 | 4.50 | 11.00 | LEU_SC1 | 9.50 | 21.50 | SER_SC1 | 4.00 | 9.00 | TYR_SC2 | 4.50 | 11.00 |
| PHE_SC3 | 4.50 | 11.00 | MET_BB | 9.00 | 22.00 | THR_BB | 9.50 | 23.00 | TYR_SC3 | 4.50 | 11.00 |
| GLY_BB | 9.50 | 23.00 | MET_SC1 | 11.50 | 22.50 | THR_SC1 | 7.00 | 17.00 | TYR_SC4 | 4.00 | 8.50 |

**Tab. S2. Parameters of the forward model to predict a density map from a Martini 3 coarse grained model.** $B_k$ parameters are expressed in Å$^2$.

| PDB id | EMDB id | res [Å] | # prot resid | # prot chains | # waters | # lipids | # ligands | box size [Å] | # waters | # lipids | # buffer ions | # atoms |
|---|---|---|---|---|---|---|---|---|---|---|---|---|
| 7p6a | 13223 | 1.90 | 540 | 5 | 76 | 0 | 0 | 122-70-48 | 10,782 | 0 | 124 | 40,805 |
| 7w9w | 32377 | 2.02 | 810 | 3 | 144 | 24 | 0 | 90-90-113 | 17,548 | 169 | 113 | 86,452 |
| 7n00 | 24095 | 2.27 | 806 | 4 | 0 | 0 | 0 | 115-115-115 | 43,754 | 0 | 260 | 143,124 |
| 7t4n | 25681 | 2.35 | 914 | 2 | 0 | 0 | 0 | 150-150-150 | 100,987 | 0 | 574 | 318,262 |
| 7b5o | 12042 | 2.50 | 651 | 3 | 73 | 0 | 1 | 120-120-120 | 50,944 | 0 | 289 | 163,693 |
| 7mjs | 23883 | 3.03 | 715 | 3 | 0 | 1 | 2 | 90-90-156 | 29,943 | 190 | 168 | 126,588 |
| 7lq6 | 23482 | 3.28 | 717 | 1 | 0 | 0 | 0 | 139-139-139 | 80,921 | 0 | 465 | 254,485 |
| 7nqk | 12528 | 3.50 | 781 | 2 | 0 | 0 | 0 | 115-115-143 | 39,764 | 328 | 222 | 175,791 |
| 6yeg | 10792 | 4.00 | 2064 | 12 | 0 | 0 | 0 | 132-132-132 | 60,592 | 0 | 496 | 212,980 |

**Tab. S3. Details of the systems used in the single-structure and ensemble refinement benchmark.** For each system, the first 8 columns report details about the deposited structure: PDB and EMDB ids, resolution, number of protein residues, number of protein chains, number of resolved water molecules, number of resolved lipids, number of ligands. The last 5 columns contain details of the system as prepared for EMMIVox simulations: box size, number of water molecules, number of lipids, number of buffer ions, and total number of atoms.

### List of Supplementary Figures

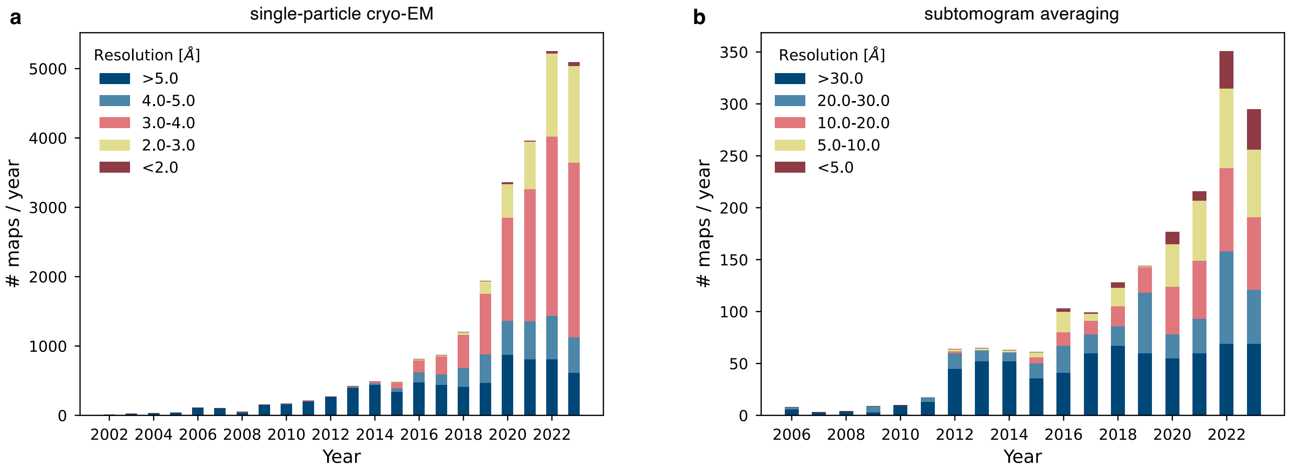


**Fig. S1. The cryo-EM and cryo-ET revolutions.** Number of (**a**) single-particle cryo-EM and (**b**) subtomogram averaging maps released per year and classified based on resolution. Data was obtained from the Electron Microscopy Data Bank (https://www.ebi.ac.uk/emdb/) on October 12th 2023.

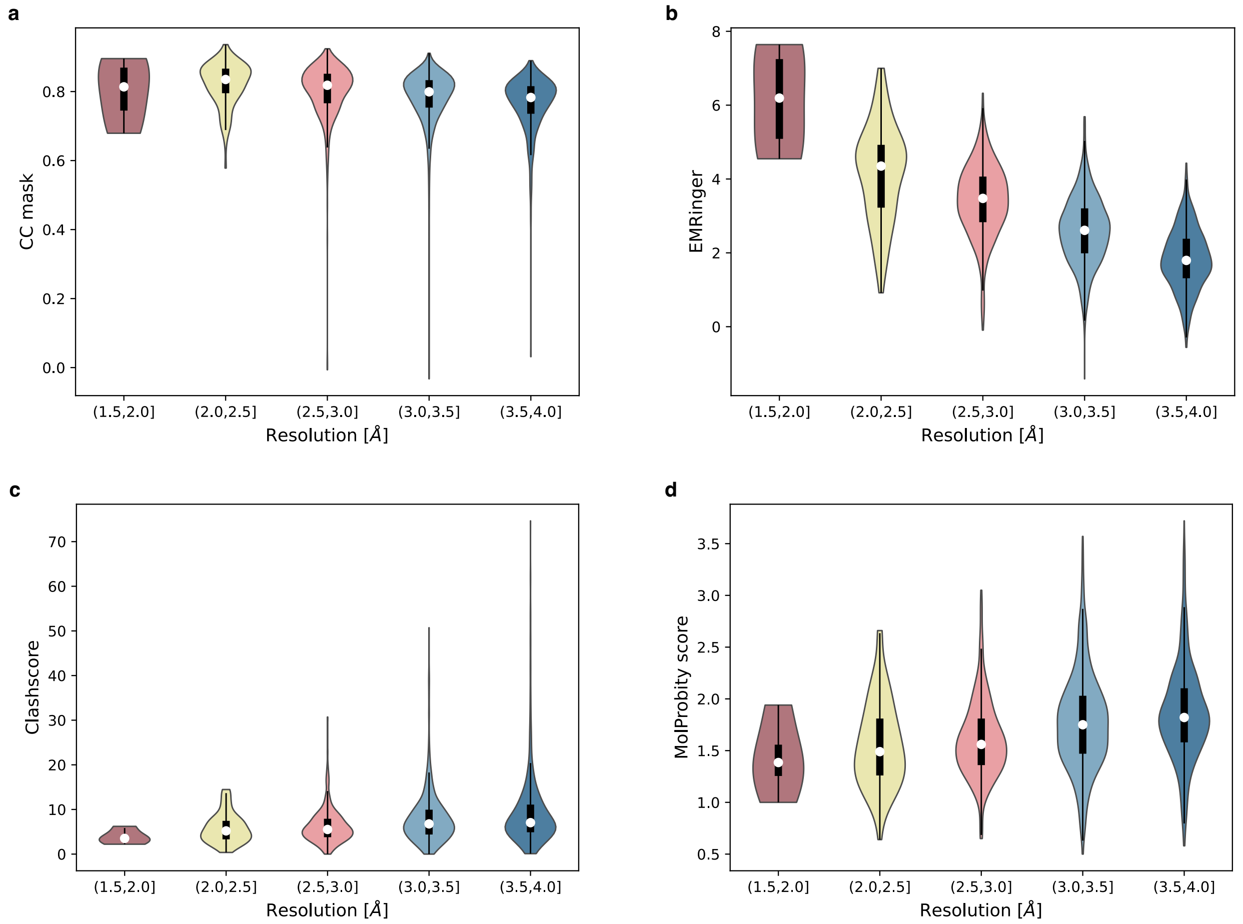


**Fig. S2. Assessment of the quality of the cryo-EM structures deposited in the PDB database.** Violin plots of (**a**) $CC_{mask}$, (**b**) EMRinger score, (**c**) clashscore, and (**d**) MolProbity score as a function of the resolution of the cryo-EM structures. For this analysis, 2476 structures determined by single-particle cryo-EM in the resolution range 1.78 Å to 4.00 Å and released between 2015 and 2022 were used. In the violin plots, the white circle corresponds to the median value, the black rectangle extends from the first to the third quantiles, and the thin black line represents the 95% confidence intervals.

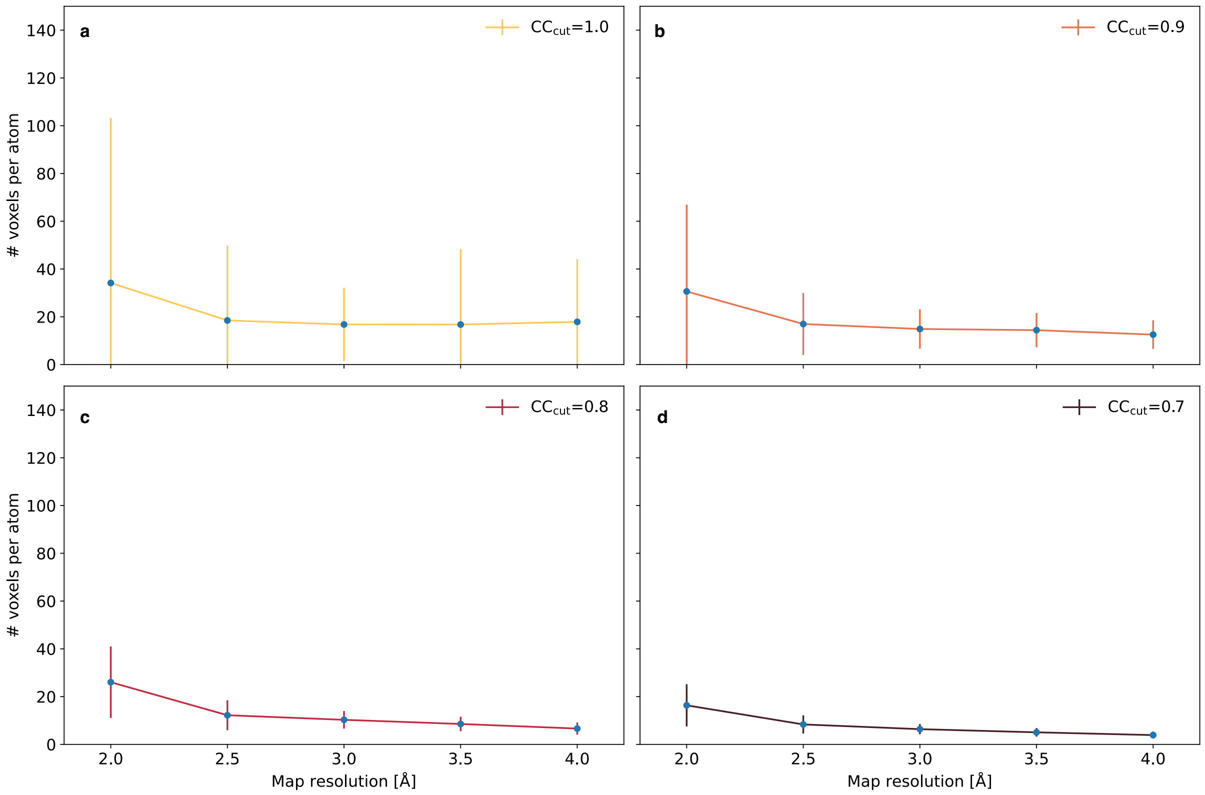


**Fig. S3. Benchmark of the approach to remove correlated voxels.** Analysis of the number of voxels per atom for 2477 structures determined by single-particle cryo-EM between 2015 and 2022 in the resolution range 1.78 Å to 4.00 Å as a function of the map resolution and voxel correlation cutoff value ($CC_{cut}$). Maps were separated into 5 resolution bins of size equal to 0.5 Å and ranging from 1.75 Å to 4 Å. The median values within the 5 bins are reported as blue points and the standard deviations as error bars. The reduction of the error bar at fixed resolution bin observed as $CC_{cut}$ is decreased indicates that upon removing correlated data the number of voxels per atom across different maps of similar resolution depends less and less on the voxel size.

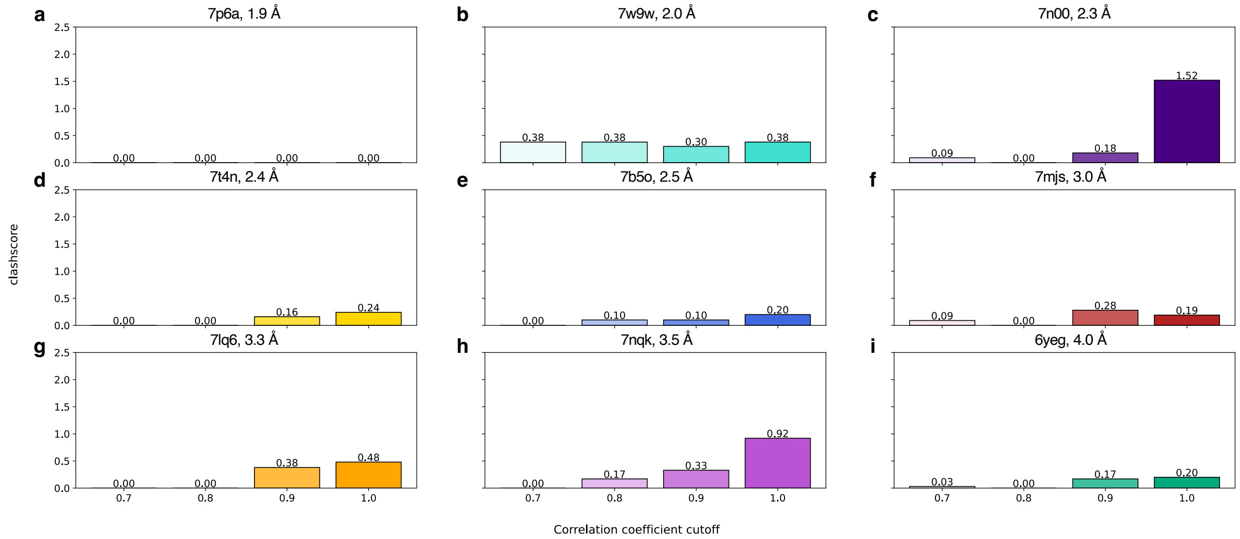


**Fig. S4. Stereochemical quality of single-structure EMMIVox models as a function of data correlation cutoff.** Comparison of clashscore for EMMIVox refined single-structure models as a function of correlation coefficient cutoff for the 9 systems in our benchmark set.

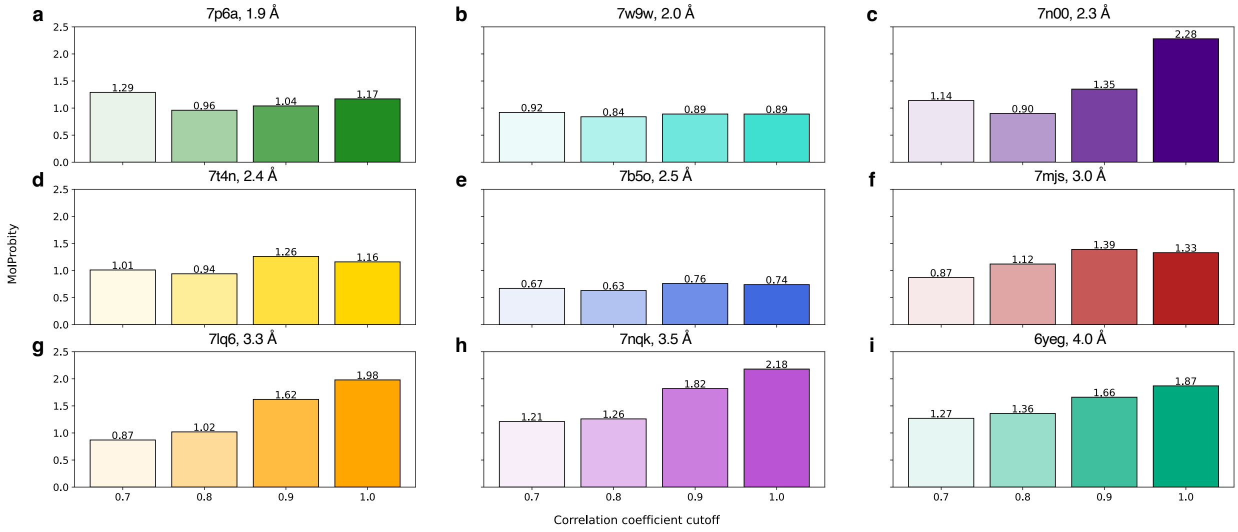


**Fig. S5. Stereochemical quality of single-structure EMMIVox models as a function of data correlation cutoff.** Comparison of MolProbity scores for EMMIVox refined single-structure models as a function of correlation coefficient cutoff for the 9 systems in our benchmark set.

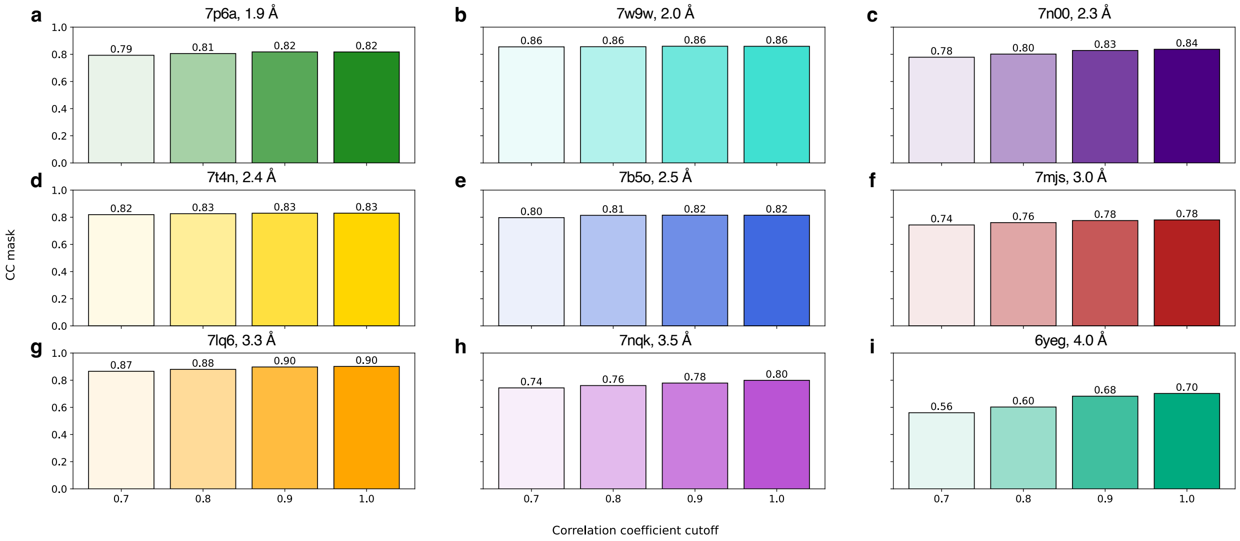


**Fig. S6. Fit-to-data of single-structure EMMIVox models as a function of data correlation cutoff.** Comparison of $CC_{mask}$ scores for EMMIVox refined single-structure models as a function of correlation coefficient cutoff for the 9 systems in our benchmark set.

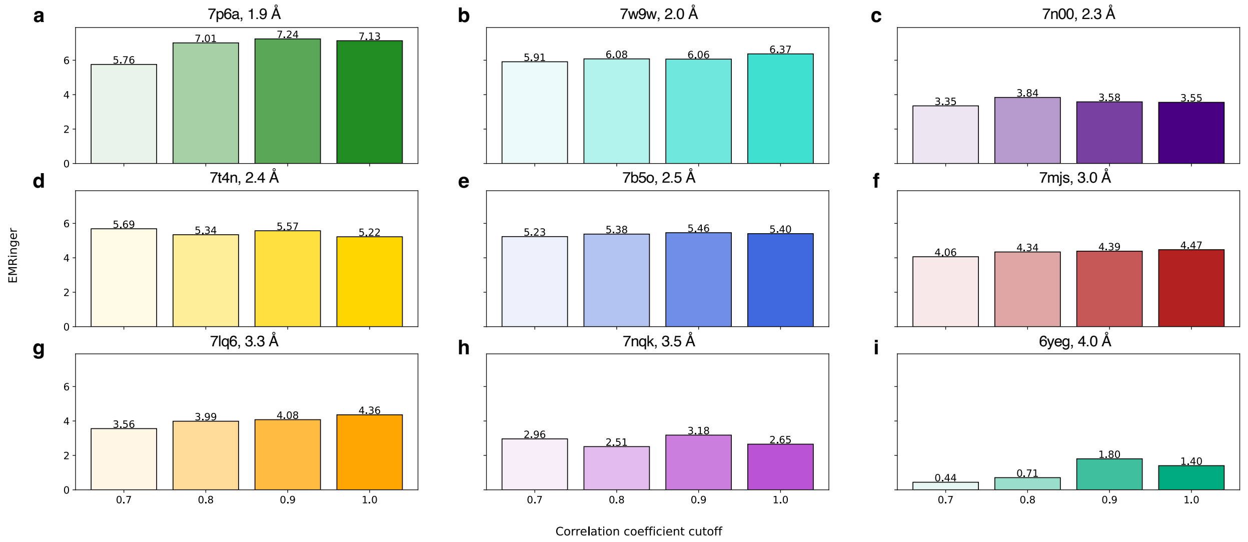


**Fig. S7. Fit-to-data of single-structure EMMIVox models as a function of data correlation cutoff.** Comparison of EMRinger scores for EMMIVox refined single-structure models as a function of correlation coefficient cutoff for the 9 systems in our benchmark set.

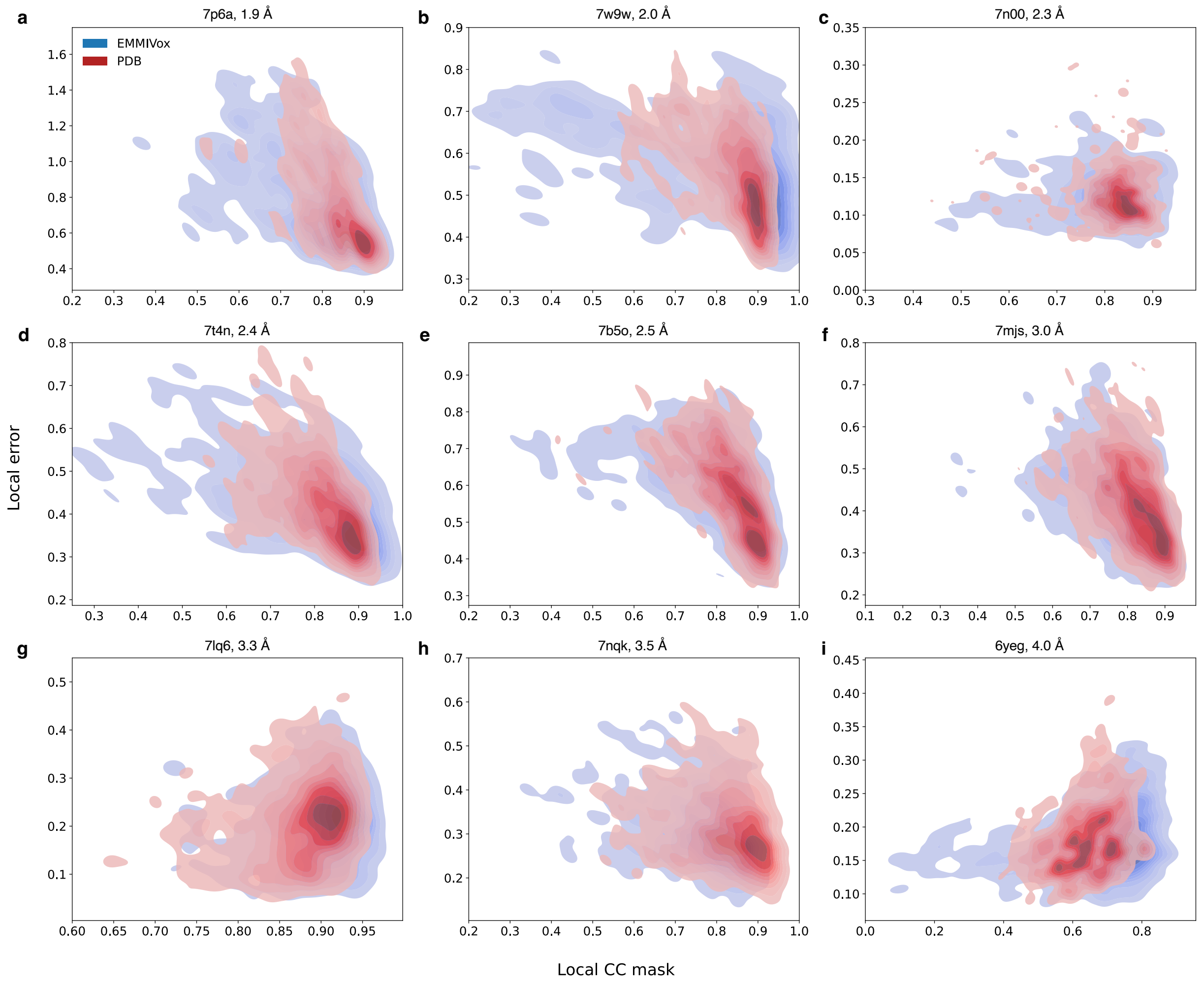


**Fig. S8. Effect of EMMIVox noise models on model fit-to-data.** Relation between individual residue fit to the experimental cryo-EM map (local $CC_{mask}$) and local error in the map in the proximity of each residue for the single-structure EMMIVox model (blue) and the deposited PDB (red) for the 9 systems in our benchmark set. The local error is calculated from the difference between the two cryo-EM half maps as the median value across all the voxels associated to a residue by Voronoi tessellation.

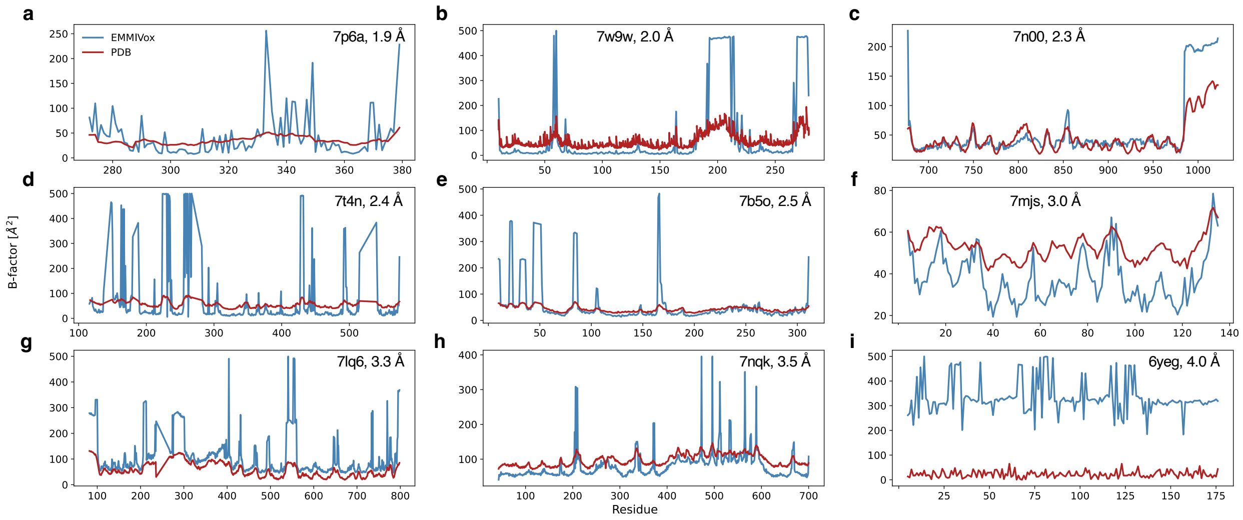


**Fig. S9. Bayesian inference of B-factors in EMMIVox single-structure refinement.** Comparison between the B-factor values of the deposited PDBs (red) and the EMMIVox single-structure models (blue) for the 9 systems in our benchmark set. Each panel compares the B-factor values for a single chain within the system and is representative of the system as a whole.

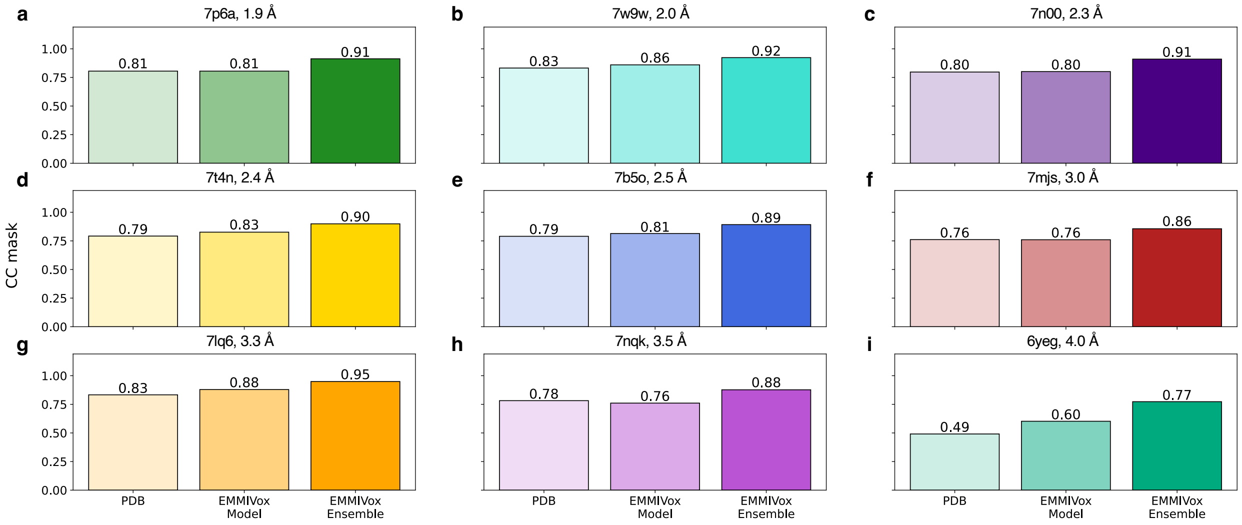


**Fig. S10. Fit-to-data of deposited PDBs, EMMIVox single-structure models and ensembles.** Comparison of the $CC_{mask}$ scores for the deposited PDB (light color), EMMIVox single-structure model (medium color) and EMMIVox ensemble (dark color).

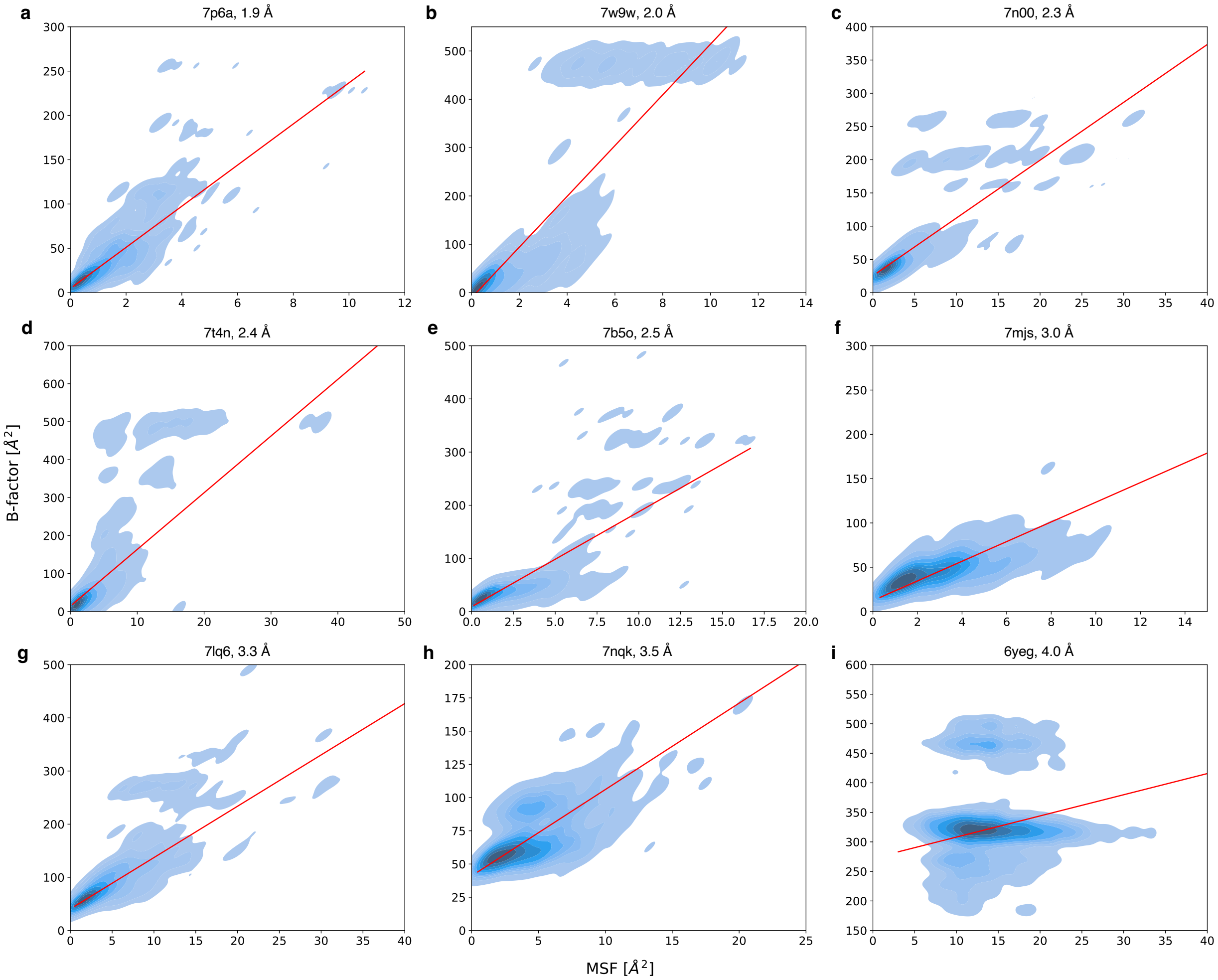


**Fig. S11. Assessment of overfitting of data noise with conformational heterogeneity in EMMIVox structural ensembles.** Relation between per residue B-factor in the single-structure EMMIVox model and residue mean square fluctuations (MSF) within the EMMIVox ensemble for the 9 systems in our benchmark set. The red line indicates a Bayesian linear fit between the two quantities.

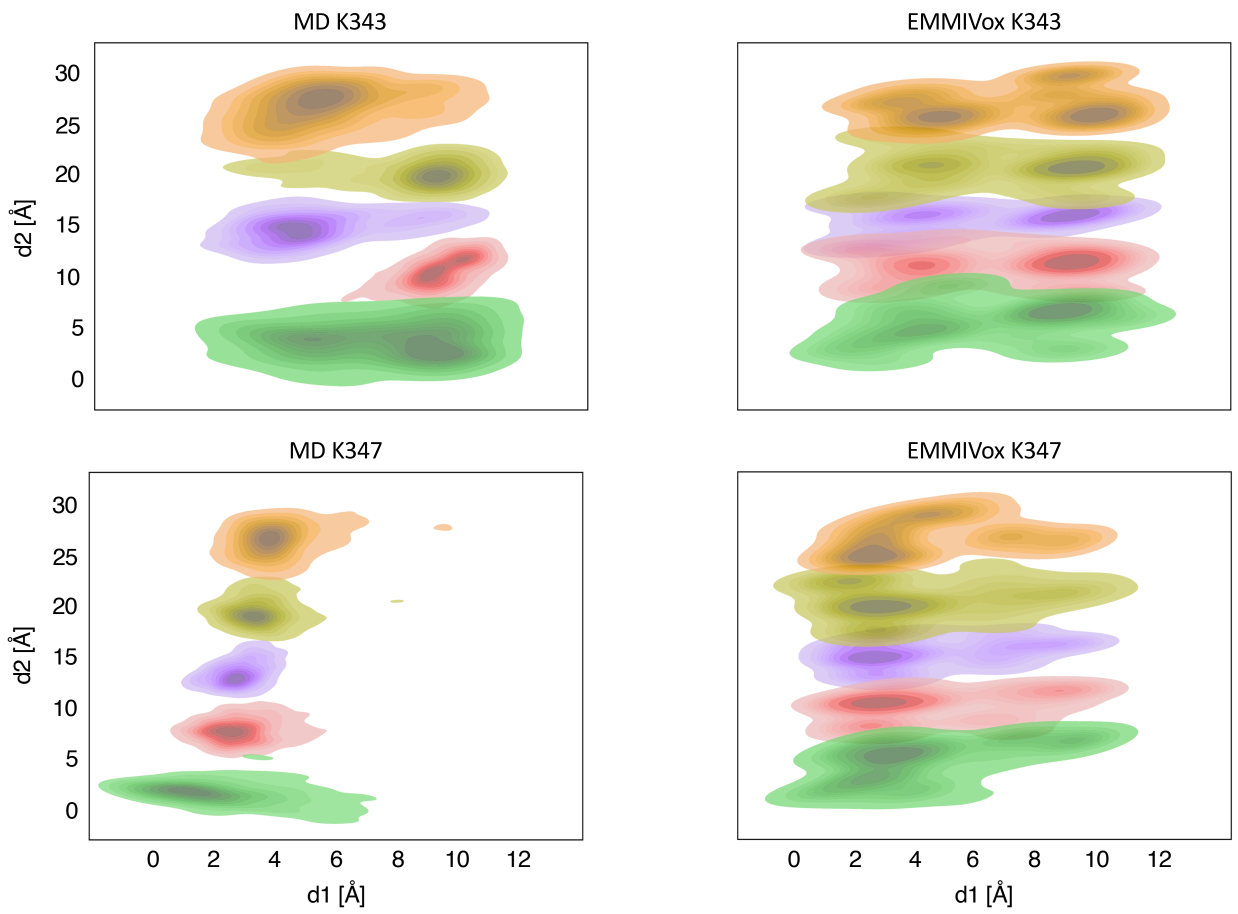


**Fig. S12. EMMIVox sampling efficiency for the case of the GPT type 1a tau filament.** Structural ensembles of residues K343 (top row) and K347 (bottom row) from standard MD simulation (left column) and EMMIVox ensembles (right column). Total simulation time for both ensembles was equal to 320 ns. In the EMMIVox ensemble, K343 occupies two distinct states with equal populations, as illustrated by the distribution of the lysine sidechain nitrogen positions projected on a plane parallel to the fibril surface. The MD ensemble for K343 shows a 2-3 split of chains occupying state 1 and state 2. Limited sampling is seen between the two states for each individual chain. K347 in the EMMIVox ensemble occupies two conformations, the major conformation found in the deposited PDB with an occupancy of 73%, and a second minor conformation, populated only by 27%. In contrast, K347 for the standard MD simulation only occupies the major conformation found in the deposited PDB and does not sample the minor conformation in the same simulation time of the EMMIVox ensemble.

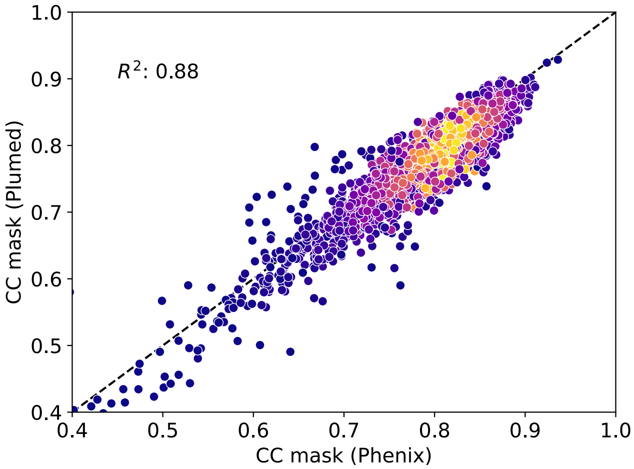


**Fig. S13. Correlation between PLUMED and Phenix $CC_{mask}$.** Analysis of the relation between $CC_{mask}$ calculated with Phenix and with a custom python script for 2477 structures determined by single-particle cryo-EM between 2015 and 2022 in the resolution range 1.78 Å to 4.00 Å. Our implementation of $CC_{mask}$ does not optimize an isotropic B-factor, unlike Phenix. Despite this difference, a strong correlation between the two $CC_{mask}$ scores is observed across the analyzed systems. Points are colored based on local density, with yellow and blue indicating high and low densities, respectively.

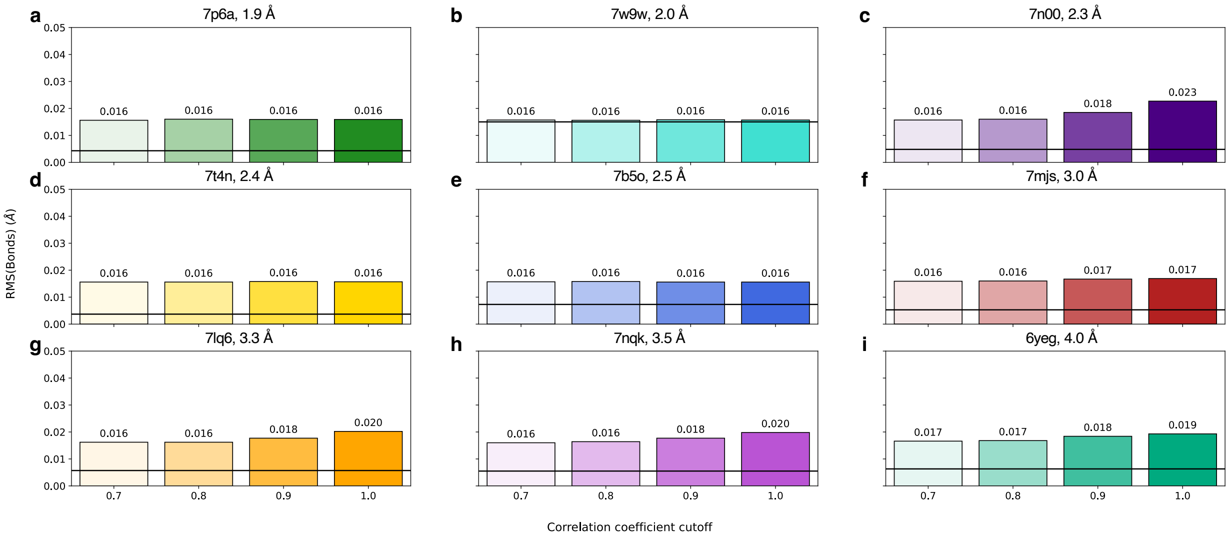


**Fig. S14. Fit-to-data of single-structure EMMIVox models as a function of data correlation cutoff.** Comparison of RMS(Bonds) values for EMMIVox refined single-structure models as a function of correlation coefficient cutoff for the 9 systems in our benchmark set. Most values fall under the 0.02 Å threshold used to define high-resolution models[1, 2]. The black line indicates the reference value in the deposited PDB.

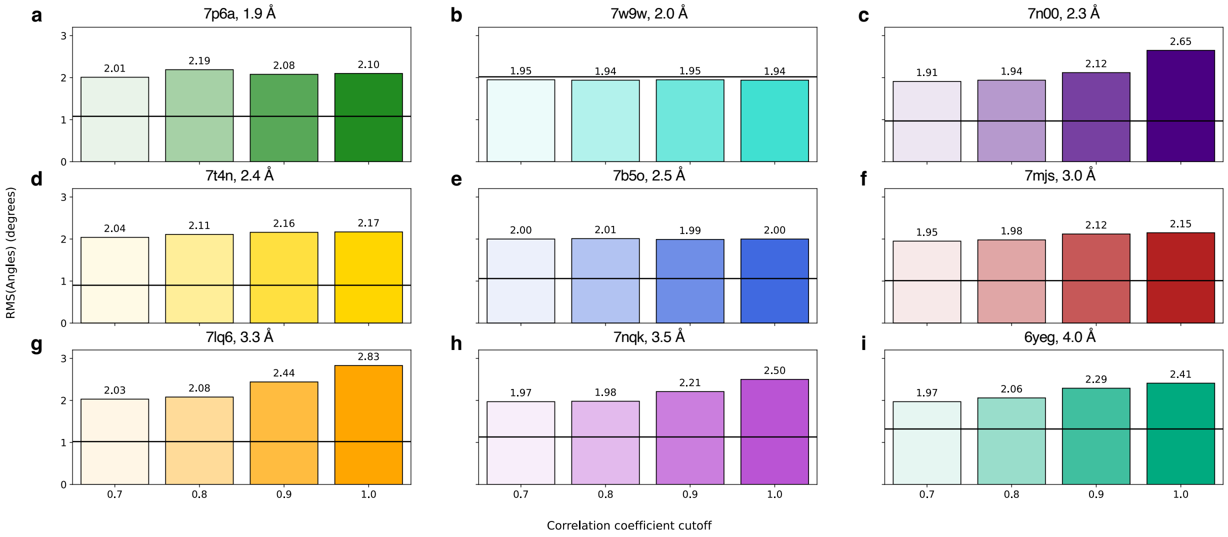


**Fig. S15. Fit-to-data of single-structure EMMIVox models as a function of data correlation cutoff.** Comparison of RMS(Angles) values for EMMIVox refined single-structure models as a function of correlation coefficient cutoff for the 9 systems in our benchmark set. Most values are within the common range of RMS(angle) values[1-3]. The black line indicates the reference value in the deposited PDB.

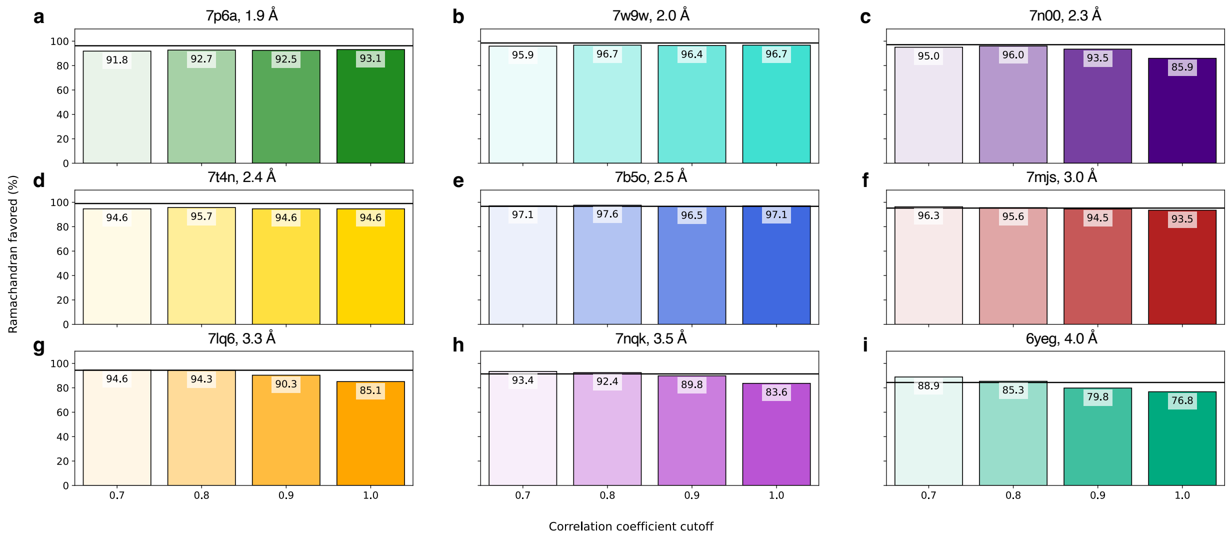


**Fig. S16. Fit-to-data of single-structure EMMIVox models as a function of data correlation cutoff.** Comparison of percentage of residues with dihedral values that fall in the favored Ramachandran region for EMMIVox refined single-structure models as a function of correlation coefficient cutoff for the 9 systems in our benchmark set. The black line indicates the reference value in the deposited PDB.

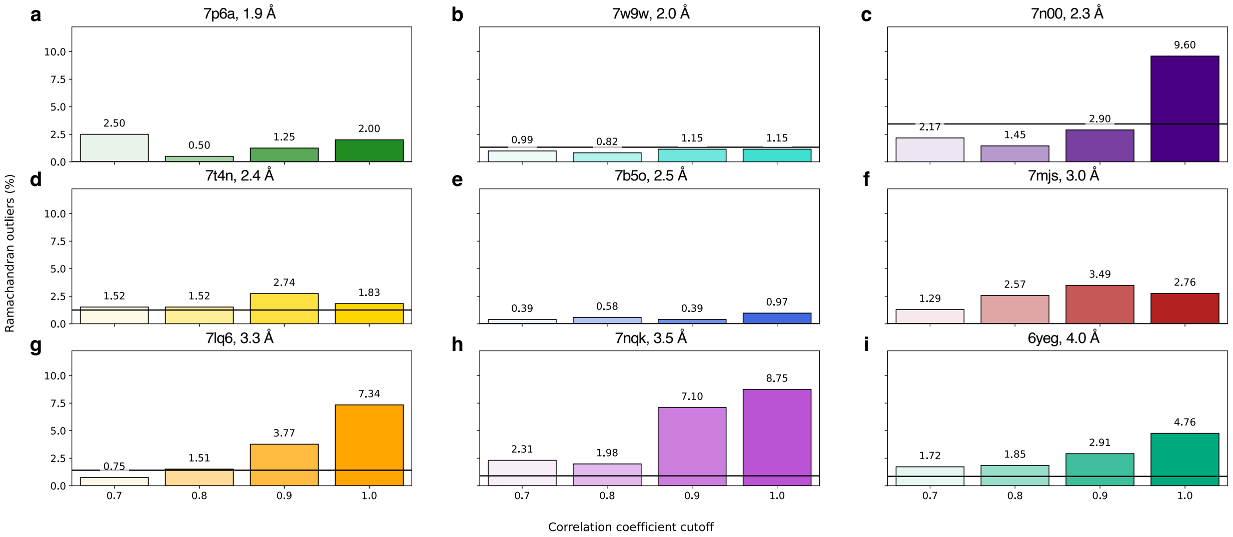


**Fig. S17. Fit-to-data of single-structure EMMIVox models as a function of data correlation cutoff.** Comparison of percentage of residues with dihedral values that fall in the non-favored Ramachandran region for EMMIVox refined single-structure models as a function of correlation coefficient cutoff for the 9 systems in our benchmark set. The black line indicates the reference value in the deposited PDB.
